# A Burst-dependent Thalamocortical Substrate for Perceptual Awareness

**DOI:** 10.1101/2023.07.13.548934

**Authors:** Christopher J. Whyte, Eli J. Müller, Jaan Aru, Matthew Larkum, Yohan John, Brandon R. Munn, James M. Shine

## Abstract

Contemporary models of perceptual awareness lack tractable neurobiological constraints. Inspired by recent cellular recordings in a mouse model of tactile threshold detection, we constructed a biophysical model of perceptual awareness that incorporated essential features of thalamocortical anatomy and cellular physiology. Our model reproduced, and mechanistically explains, the key *in vivo* neural and behavioural signatures of perceptual awareness in the mouse model, as well as the response to a set of causal perturbations. We generalised the same model (with identical parameters) to a more complex task – visual rivalry – and found that the same thalamic-mediated mechanism of perceptual awareness determined perceptual dominance. This led to the generation of a set of novel, and directly testable, electrophysiological predictions. Analyses of the model based on dynamical systems theory show that perceptual awareness in simulations of both threshold detection and visual rivalry arises from the emergent systems-level dynamics of thalamocortical loops.

## Introduction

The study of perceptual awareness – the process of gaining conscious access to perceptual content – in human participants (e.g. Overgaard, 2015; Pitts et al., 2014; Sergent et al., 2005) and animal models (e.g. Ciceri et al., 2024; Gale et al., 2024; Oude Lohuis et al., 2022; Palagina et al., 2017) have opposing but complementary limitations. Human participants can rapidly learn complex tasks that isolate and control for key psychological constructs, however the high-resolution (i.e., cell specific) recordings and precise causal manipulations (e.g., optogenetic and pharmacological) that are needed to make effective inferences about the neural basis of behaviour are exceedingly difficult and often impossible to obtain. At the same time, animal models, and transgenic mouse models in particular, allow for an astonishing degree of experimental precision in the recording and causal manipulation of neural activity. Animal models are, however, highly limited in the range and complexity of the tasks they can perform, restricting the type of psychological inferences that can be drawn. Both fields contain crucial pieces of the puzzle for understanding perceptual awareness, however the links between the two are limited at best. Effective progress, therefore, hinges on our ability to create empirically tractable tethers between the behavioural signatures of perceptual awareness studied in humans and the fine-grained neurobiological mechanisms studied in animal models (He, 2023).

Recent work in a mouse model of perception has identified a key thalamocortical circuit connecting thick-tufted layer 5 pyramidal-tract (L5PT) neurons and matrix thalamic cells as playing a causal role in the threshold for perceptual awareness (Aru et al., 2019, 2020; Bachmann et al., 2020; Takahashi et al., 2016, 2020). Specifically, based on a range of cellular recordings and causal perturbations, it has been shown that matrix-thalamus-mediated coupling of apical dendrite and somatic compartments in L5PT cells leads to a burst-firing state that is a reliable signature of perceptual awareness of a near-threshold tactile stimulus (Takahashi et al., 2016, 2020). However, the simplicity of the threshold detection task and species-specific differences in neural architecture means that it is not clear whether the mechanisms of perceptual awareness characterised in the mouse model will generalise beyond the whisker detection task to the more complex paradigms typically studied in human participants.

Here, we use biophysical modelling to bridge the gap between the thalamocortical circuit identified in the mouse model of perception (Aru et al., 2020; Takahashi et al., 2016a, 2020) and the behavioural signatures of perceptual awareness studied in human psychophysics. Specifically, we built a thalamocortical spiking neural network model that explains the full suite of behavioural and neural findings in the mouse model of tactile threshold detection. Given the ubiquity of the thalamocortical circuit architecture across sensory modalities, we (Aru et al., 2020; Bachmann et al., 2020; Whyte et al., 2024), along with others (Marvan et al., 2021; Phillips et al., 2016), have proposed that reverberant bursting activity in L5PT – matrix thalamus loops may be a necessary component part in a domain general mechanism of perceptual awareness. A key test of this hypothesis is whether this same circuit architecture can explain psychophysical principles known to govern perceptual awareness in more complex paradigms and in other sensory modalities.

To test this hypothesis *in silico*, we leveraged the same model with identical parameters to simulate both tactile threshold detection and visual rivalry (which we use as a catch all term for binocular rivalry and related bistable perception paradigms). Visual rivalry is a complex but highly psychophysically-constrained phenomenon whereby visual perception stochastically switches between stimulus percepts that differ only in terms of their perceptual content (Alais & Blake, 2005; Blake & Tong, 2008; Carmel et al., 2010; Tong et al., 2006). Visual rivalry provides a means to dissociate the neural mechanisms of subjective perception from the correlates of physical stimulation. Crucially, variants of visual rivalry (i.e., plaid rivalry; Hupé & Rubin, 2003) can now be studied in mouse models (Bogatova et al., 2022; Palagina et al., 2017) as well in human and non-human primates, allowing us to test our model against existing psychophysical findings in primates and to make predictions for what should be observed in the mouse model before data has been collected – a central component in the evolving dialogue between theory and experiment. In addition, although our model is consistent with the psychophysical predictions of previous models of visual rivalry (e.g., Grossberg et al., 2008; Laing et al., 2010; Laing & Chow, 2002; Safavi & Dayan, 2022; Shpiro et al., 2007, 2009; Wilson, 2003, 2007, 2017), our approach has an unique level of neurobiological specificity that allows us to generate cellular level predictions about the neural underpinnings of perceptual awareness in a language that is applicable to the causal methods used by modern systems neuroscientists.

## Results

### A spiking corticothalamic model recreates key features of cellular physiology

The dynamical elements of the model were inspired by recent empirical observations, and consist of three classes of neurons (Fig 1 A-B) – L5PT cells (blue), fast spiking interneurons (basket cells, gold), and diffuse projecting matrix thalamocortical cells (purple) – each of which are modelled using biophysically plausible spiking neurons (Izhikevich, 2003, 2004, 2006; Naud & Sprekeler, 2018) that were coupled through conductance-based synapses. By building the model from these circuit elements, we ensured that the emergent dynamics of the population recapitulate known signatures of cell-type-specific firing patterns, thus retaining the capacity to translate insights between computational modellers and cellular physiologists.

**Fig 1.**
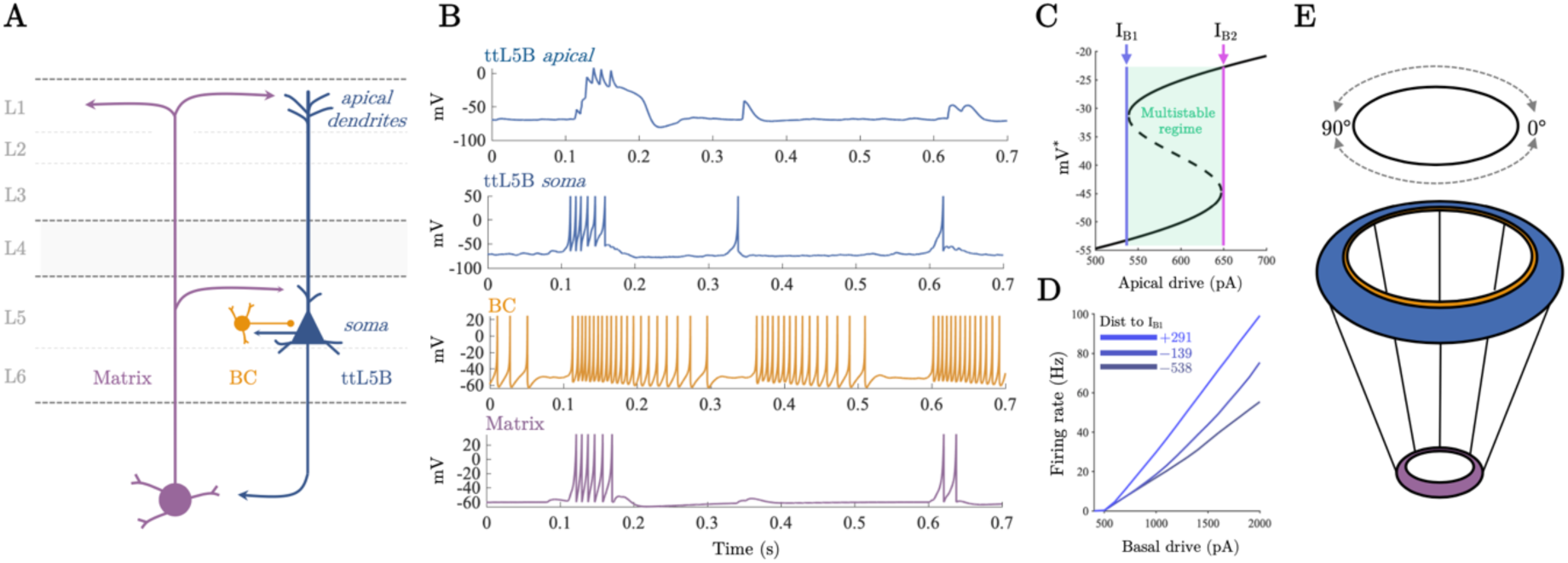
Thalamocortical model of perceptual awareness. A) Idealised anatomy of the model thalamocortical loop connecting higher-order matrix thalamus and L5_PT_ neurons. B) Single neuron dynamics of example neurons in the thalamocortical network for each class of cell when driven with 600 [Hz] of independent background drive to the somatic compartment of every neuron and 50 [Hz] to apical compartment of L5_PT_ cells. C) Bifurcation diagram of the L5_PT_ apical compartment. I_B1_ denotes a saddle node bifurcation generating a stable plateau potential which coexists with the resting state of the apical compartment until the model passes through a second saddle node bifurcation at I_B2_ at which point the resting state vanishes and the stable plateau potential becomes globally attracting. D) Somatic firing rate of the novel dual compartment L5_PT_ model as a function of basal (i.e. somatic compartment) drive and apical drive (measured in terms of the distance to I_B1_). In line with empirical data (Larkum, 2004), apical drive increases the gain of the somatic compartment. E) Model thalamocortical ring architecture. L5_PT_ cells and basket cells were placed on a cortical ring at evenly space intervals.

To model the non-linear bursting behaviour of L5PT cells (which has been linked to perceptual awareness; Larkum, 2004, 2022; Larkum et al., 2009), we created a novel dual-compartment model with active apical dendrites that captures the essential features of the cells’ physiology. Empirical recordings have shown that L5PT cells switch from regular spiking to bursting when they receive near-simultaneous input to both their apical (top) and basal (bottom) dendrites (Larkum, 2004, 2022; Fig 1B). As such, our model includes a somatic compartment, described by an Izhikevich adaptive quadratic integrate and fire neuron (Izhikevich, 2006; Munn, Müller, Aru, et al., 2023; Munn, Müller, Medel, et al., 2023) and an apical compartment, described by a non-linear model of the Ca^2+^ plateau potential (Naud et al., 2014; Naud & Sprekeler, 2018). To recapitulate known physiology, we coupled the compartments such that sodium spikes in the somatic compartment back propagate to the apical compartment; in turn if a Ca^2+^ plateau potential is triggered in the apical compartment the somatic compartment’s behaviour (probabilistically) switches from regular spiking to bursting (implemented by switching the reset conditions of the somatic compartment). The amount of current entering the apical compartment controls this switching process. With sufficiently high current the apical compartment passes through a saddle node bifurcation (IB1; Fig 1C) and a stable Ca^2+^ plateau potential coexists with the resting state of the compartment. Further increases in current cause the cell’s resting state to disappear (by passing the cell through a second saddle node bifurcation at IB2; Fig 1C), making the plateau potential globally attracting (supplementary materials S1A- D).

Based on the finding that communication between the soma and apical dendrites of L5PT cells depends upon depolarising input from the matrix thalamus to the “apical coupling zone” of L5PT cells (Suzuki & Larkum, 2020), we made the probability of successful propagation between compartments proportional to the amplitude of thalamic conductances. In line with empirical findings and previous modelling (Larkum, 2004; Shai et al., 2015), Ca^2+^ plateau potentials in the apical compartment controlled the gain of the somatic compartment’s firing rate curve by increasing the amount of time the somatic compartment spent in a bursting rather than a regular spiking parameter regime (Fig 1D).

In line with previous spiking neural network models of early sensory cortex (Laing et al., 2010; Laing & Chow, 2002; Wang et al., 2020; Zerlaut et al., 2018), we embedded the cortical neurons in a one-dimensional ring architecture (90 pairs of L5PT excitatory and fast-spiking inhibitory interneurons; Fig 1E). Each point on the ring represents an orientation preference, with one full rotation around the ring corresponding to a 180^°^ visual rotation – this provides each neuron with a 2^°^ difference in orientation preference, relative to its neighbours. The cortical ring was coupled to a thalamic ring with a 9:1 ratio (to approximately reflect the cortico-thalamic ratio in mammals), which then projected back up to the apical dendrites of the same 9 cortical neurons, representing the diffuse projections of higher-order thalamus onto the apical dendrites of L5PT neurons in layer 1 (Mease & Gonzalez, 2021; Shepherd & Yamawaki, 2021). Cortical coupling was modelled with a spatial decay, with long range inhibitory coupling and comparatively local excitatory coupling (i.e., centre-surround ‘Mexican-hat’ connectivity).

When driven solely by baseline input, the model emitted irregular spikes interspersed with sparse spatially localised bursts mediated by depolarising input from the thalamus which allowed L5PT somatic spikes to back-propagate initiating Ca^2+^ plateau potentials in the apical dendrites which in turn initiated transient burst spiking in the somatic compartment (Fig 1C). Cortical spiking activity was highly irregular (mean inter-spike interval coefficient of variation 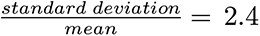) characteristic of a waking state (Destexhe, 2009). For more details on the model architecture and analysis of the dynamics see materials and methods.

#### The thalamocortical model reproduces empirical signatures of threshold detection

We first set out to reproduce the results of the whisker-based tactile detection paradigm employed by Takahashi and colleagues (2016, 2020), who trained mice to report a mechanical deflection of a whisker over a range of deflection intensities (Fig 2A) whilst recording L5PT activity in barrel cortex from the apical dendrites via fast scanning two-photon Ca^2+^ imaging, and somatic activity via juxtacellular electrodes. The original study found that bursting activity in the soma of L5PT cells, generated by Ca^2+^ plateau potentials in the apical dendrites, distinguished hits and false alarms from misses and correct rejections. Importantly, they were able to establish causality through a series of perturbation experiments (Fig 2B). Optogenetic excitation of the apical dendrites reduced the animal’s threshold for awareness increasing both hits and false-alarms (Fig 2C). In turn, pharmocological inhibition of the apical dendites and POm (a matrix-rich higher-order thalamic nucleus with closed loop connections to barrel cortex; Mease & Gonzalez, 2021) increased the animals perceptual threshold (Fig 2D-E).

**Fig 2.**
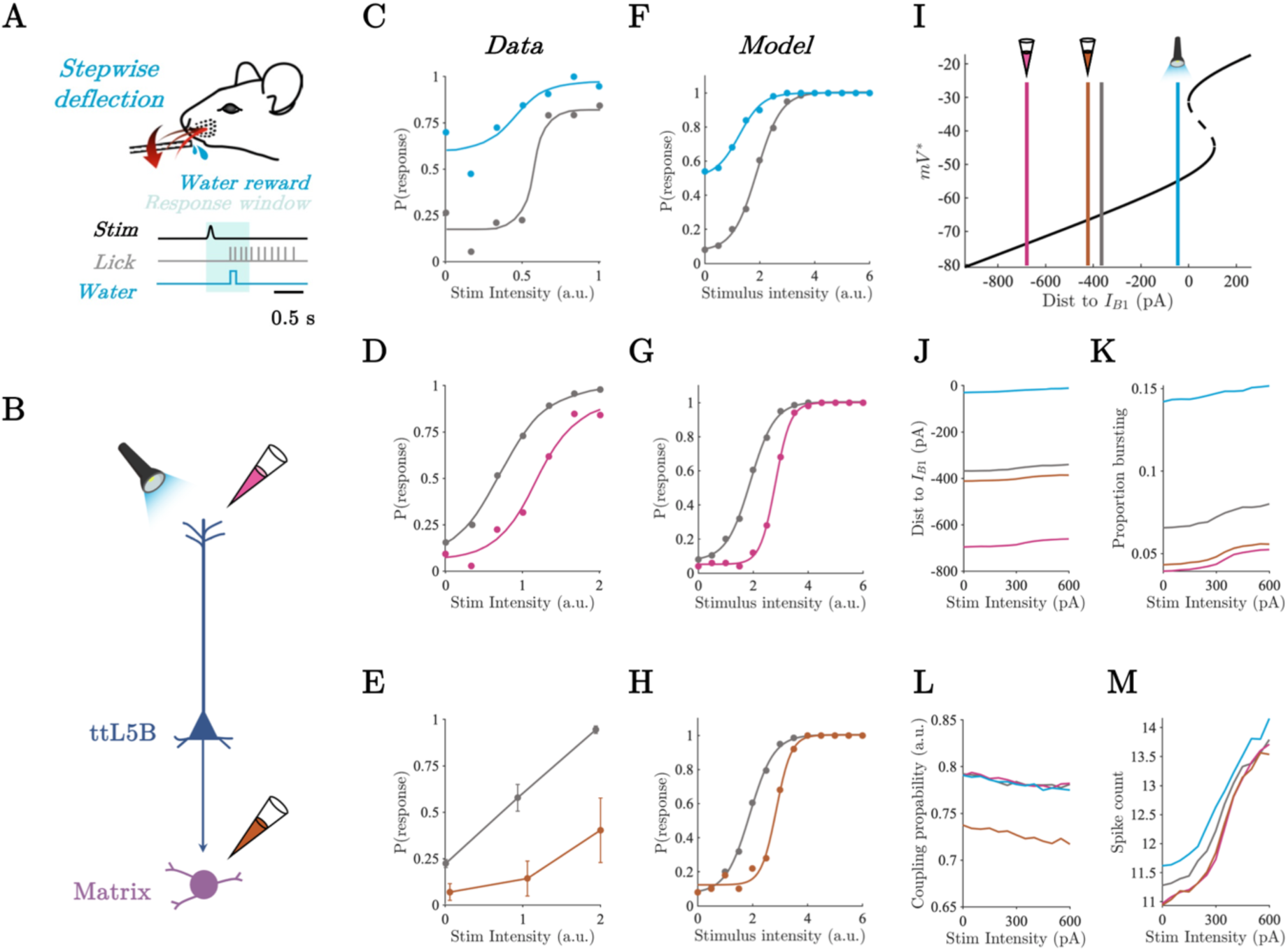
Apical compartment distance to bifurcation and thalamic-gating explains shifts in perceptual threshold. A) Whisker deflection paradigm modified from Takahashi et al, (2020). B) Representation of causal perturbations to L5_PT_ thalamocortical circuit including optogenetic excitation of the apical dendrites (light blue), pharmacological inhibition of the apical dendrites (pink), and pharmacological inhibition of POm (orange). C-D) Psychometric function of animals performing a whisker deflection task for the control condition (grey), optogenetic excitation of apical dendrites (blue), and pharmacological inhibition of apical dendrites (pink). Modified from Takahashi et al, (2016) and Takahashi et al, (2020). E) Response probability as a function of whisker deflection intensity for control (grey) and pharmacological inhibition of POm (orange). Modified from Takahashi et al., (2020). F-H) Model response probability as a function of stimulus intensity across simulated causal perturbations. Colours same as above. I) Apical compartment bifurcation diagram showing the average distance to B_1_ in post stimulus period averaged across stimulus intensities. J) Average distance to B_1_ across stimulus intensities and simulated causal perturbations in the post stimulus period. K) Proportion of population in bursting regime across stimulus intensities and simulated causal perturbations in the post stimulus period. L) Average inter-compartment coupling probability across stimulus intensities and simulated causal perturbations in the post stimulus period. M) Average spike count across stimulus intensities and simulated causal perturbations in the post stimulus period.

To model the perceptual discrimination process underlying threshold detection – discriminating the presence of a weak stimulus against a noisy background – all neurons received a constant background drive consisting of independent Poisson spike trains whilst an arbitrary cortical neuron was pulsed by a current of constant width and variable amplitude that we weighted by a spatial Gaussian to mimic the selectivity of neurons in early sensory cortex. We operationalised perceptual awareness in the threshold detection simulations in terms of what an upstream ideal observer could readout from the population by computing whether trial-by-trial spike counts in the 1000 ms post stimulus window exceeded an optimal criterion (i.e. the criterion that best minimises misses and false alarms across stimulus intensities). We counted the model as having made a response whenever the spike count exceeded the optimal criterion. Psychometric functions were then fit to the model’ responses (for details see materials and methods). Qualitatively identical results were obtained using neurometric functions which summed over all criterion values (see supplementary material S2; Britten et al., 1992).

The model responses to simulated whisker deflections recapitulated the empirically observed sigmoid-like relationship between the intensity of the simulation and the model’s response probability (Fig 2F-H). In addition, perturbations to the model designed to replicate optogenetic excitation and pharmacological inhibition (Fig 2A) qualitatively reproduced the empirically observed shifts in response probability across perturbation types (Fig 2C-H). For each type of perturbation we ran the simulation over a range of perturbation magnitudes to ensure the reliability of the effect see supplementary material S2. For brevity, we only show the results for 300 [pA] perturbations in the main text. In addition, we note that the saturating detection probability values in the model are due to an absense of behavioural stochasticity resulting from extranious factors such as decision noise which are inherent to empirical data.

A key benefit of biophysical modelling is the capacity to mechanistically probe the model and determine how the empirical observations may have emerged from the underlying circuit dynamics. To this end, we used tools from dynamical systems theory to interrogate the cell-type-specific dynamics underlying the behaviour of the model including the distance to bifurcation (IB1) in the apical compartment, the parameter regime of the somatic compartment (i.e. regular spiking or bursting), and the inter-compartment coupling probability. Excitation of the apical dendrites (blue) reduced the average distance to bifurcation (here defined as the distance to IB1) in the apical compartment across the network in the 1000 [ms] period post stimulus onset (Fig 2I- J). This increased the proportion of time each somatic compartment spent in the bursting regime (Fig 2K), which in turn increased the average spike count of L5PT cells across stimulus intensities resulting in reduction of the model’s perceptual threshold (Fig 2F & 2M). Conversely, inhibition of the apical dendrites (pink) and the thalamus (orange) both resulted in an increase in the perceptual threshold (Fig 2G- H).

Importantly, however, the mechanisms underlying the increase in the perceptual threshold differed across apical dendrite and thalamic inhibitory perturbations. Inhibition of the apical dendrites increased the average distance to bifurcation at B1 in the apical compartment across the network (Fig 2I-J). In contrast, inhibition of the thalamus resulted in a comparatively minor reduction in the distance to bifurcation in the apical dendrites (Fig 2I-J) but reduced the thalamus mediated inter-compartment coupling (Fig 2L) thereby reducing the probability that a back propagating action potential could reach the apical compartment, and the probability with which a plateau potential could switch the regime of the somatic compartment from regular spiking to bursting. Together this resulted in a similar reduction in the proportion of cells in the bursting regime for both thalamic and apical dendrite inhibition, and likewise, a similar reduction in the average stimulus evoked spike count explaining the comparable increase in perceptual thresholds (Fig 2M).

### Thalamocortical spiking model generalises to visual rivalry

We next sought to generalise our thalamocortical model of perceptual awareness to visual rivalry formalising and interrogating the hypothesis that the role played by pulvinar – L5PT loops in visual cortex is analogous to the role played by POm – L5PT loops in barrel cortex. To simulate visual rivalry we drove the model with input representing orthogonal gratings presented to each eye, typical of standard binocular rivalry experiments (Logothetis & Leopold, 1996; Tong et al., 2006), targeting the soma (i.e., basal dendrites) of L5PT cells on opposite sides of the ring with orthogonal orientation preferences (Fig 3A).

**Fig 3.**
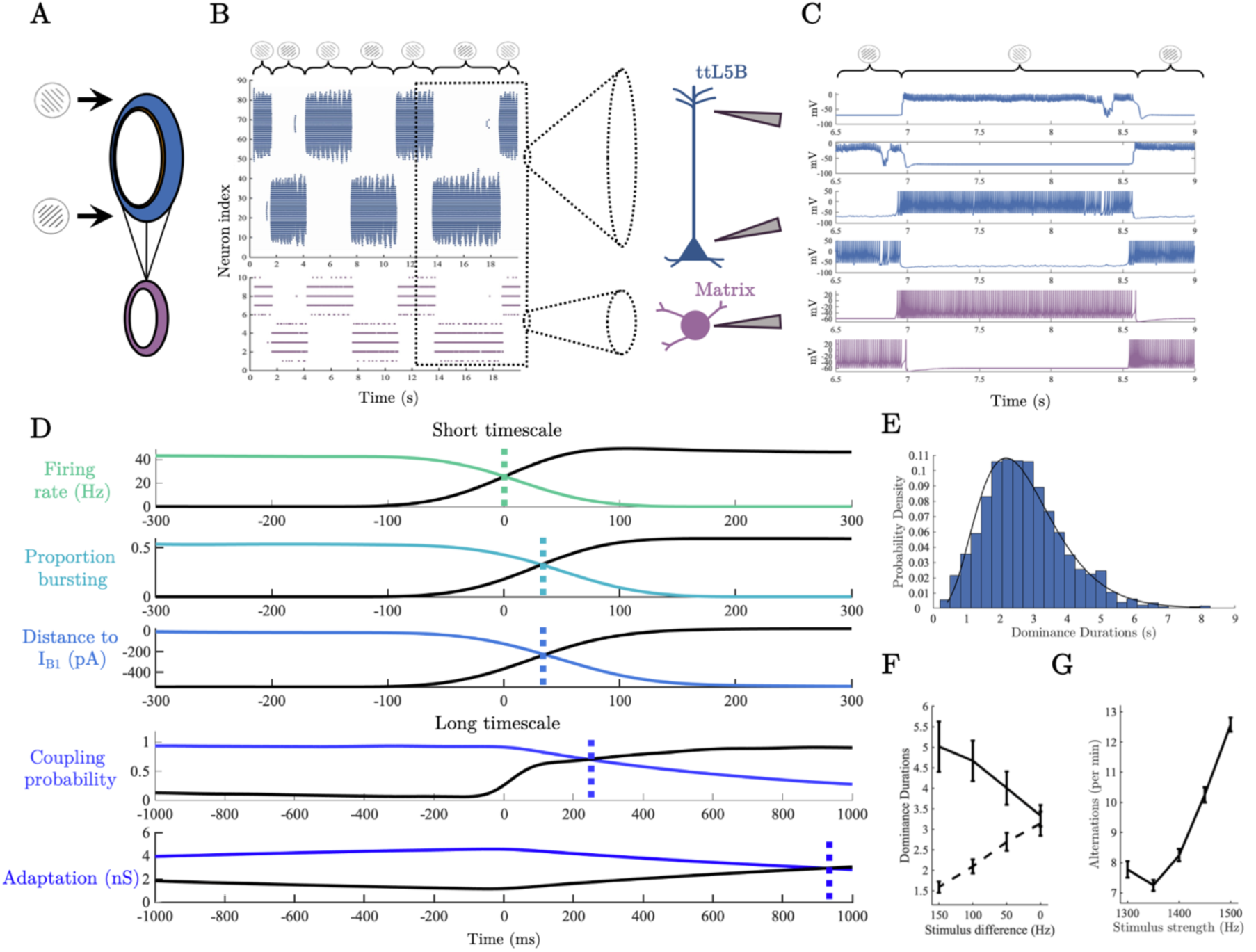
A thalamocortical cascade underlies perceptual switches. A) Thalamocortical ring architecture driven by input representing orthogonal gratings. B) Raster plots of L5_PT_ soma (blue) and matrix thalamus population (purple) during rivalry. C) Example single neuron spiking activity around a perceptual switch – dotted lines denote the crossing in the population averaged time series. D) Population averaged neuronal variables centred on a perceptual switch. E) Histogram of dominance durations, black line shows the fit of a Gamma distribution with parameters estimated via MLE (𝛼 = 4.85, 𝜃 = 0.56). F) Simulation confirming Levelt’s second proposition. Dashed line shows the dominance duration of the population receiving the decreasing external drive, solid line shows dominance duration of population receiving a fixed drive. G) Simulation of Levelt’s fourth proposition. Notably, for low stimulus strengths our model, along with a number of meanfield models of visual rivalry (Shpiro et al., 2007), predicts a deviation from Levelt’s fourth proposition.

Due to the fact that inhibitory connectivity is broader than excitatory connectivity, delivering external drive to opposite sides of the ring shifts the model into a winner-take-all regime with burst-dependent persistent states on either side of the ring competing to inhibit one another. Importantly, through the accumulation of slow a hyperpolarising adaptation current in the somatic compartment of each L5PT cell (representing slow Ca^2+^ mediated K^+^ currents; McCormick & Williamson, 1989; Wilson & Cowan, 2021), burst-dependent persistent states are only transiently stable leading to stochastic switches between persistent states characteristic of binocular rivalry (Fig 3B-C). We operationalised perceptual dominance (i.e., awareness of a percept at the exclusion of the other) in terms of the difference in average firing rate between the persistent ‘bumps’ of L5PT cell activity on opposite sides of the ring. In line with common practice (e.g. Li et al., 2017) in models of rivalry a population was counted as dominant when it had a firing rate 5 [Hz] > than its competitor and lasted for longer than 250 [ms] (results were robust across a large range of threshold values). The initial competition for perceptual dominance was determined by which population of L5PT cells first established a recurrent loop with matrix-thalamus. This interaction allowed the recurrently connected population of cells to enter a bursting regime, at which point the population had sufficient activity to maintain a persistent state and inhibit the competing population into silence. In previous work we have argued that this emergent property of corticothalamic loops is a good candidate for a cellular-level correlate of perceptual awareness (Aru et al., 2020; Whyte et al., 2024).

Crucially, our model provides a cellular-level explanation of spontaneous rivalry- induced perceptual switches in terms of these same mechanisms. As the reliable initiation of Ca^2+^ plateau potentials in the model depends upon back-propagating action potentials, perceptual switches are always preceded by regular spikes in the suppressed population. Preceding each switch, the accumulation of adaptation reduces the firing rate of the dominant population to a sufficient level that the suppressed population can escape from inhibition and emit a brief series of regular spikes that transition into bursts as thalamus is recruited and Ca^2+^ plateau potentials are initiated (Fig 3C), eventually gaining sufficient excitatory activity to inhibit the previously dominant population into silence via recurrent interactions with the basket cell population. At the population level (Fig 3D), this cascade of neuronal events is characterised by an initial ramping in the somatic firing rate of the suppressed population, followed by the inter-compartment coupling probability, the proportion of the population bursting, and finally by the average distance to bifurcation in the apical compartment. Once dominant, approximately half the population is in a bursting regime at each point in time (0.5424 mean ± 0.058 standard deviation), and the average distance to bifurcation in the apical compartment fluctuates around zero (- 11.0185 ± 48.067 [pA]), with approximately a third (0.3669 ± 0.097) of the population located above the critical boundary (IB1) at each point in time. In addition, the average inter-compartment coupling probability is (0.8845 ± 0.102) allowing reliable communication between compartments. Once silenced, adaptation in the previously dominant population decays back to baseline levels and accumulates in the previously suppressed, now dominant, population. In this way, the competitive organisation of the cortico-cortical and corticothalamic loop provides a natural “flip-flop” switch that stochastically alternates between dominant percepts.

As cortical pyramidal cells are known to display considerable differences in spiking behaviour across species (Kalmbach et al., 2021) we swept the parameters of our model L5PT cell responsible for the bursting behaviour to ensure that visual rivalry (quantified by average dominance duration) is stable across a wide range of parameters. In favour of the robustness of the model, dominance durations in an empirically plausible range (i.e. with a period on the order of seconds) occurred across a wide range of parameter values (supplementary material S3).

In close agreement with psychophysical data, switches between dominance and suppression had a right-skewed distribution of dominance durations (Brascamp et al., 2015; Levelt, 1967; Logothetis & Leopold, 1996; Fig 3E). Across stimulus drive conditions (1300 – 1500 [Hz]), comparison of negative log likelihoods showed that the distribution of dominance durations was best fit by a Gamma distribution (ℓ = 2.81 × 10^−3^), compared to lognormal (ℓ = 2.88 × 10^−3^) or normal distributions (ℓ = 2.88 × 10^−3^). Because of the large size of the network and the discontinuity in the somatic compartment we could not use analytic methods to interrogate the structure of the dynamical system underlying the stochastic oscillations. Instead we used a heuristic line of argument (see Strogatz, 2018, Ch.8) combined with simulations in the absence of noise to confirm that the dynamical regime underlying these stochastic oscillations likely consists of noisy excursions around a stable closed orbit (i.e. a stable limit cycle; supplementary material S4).

### Thalamocortical spiking model conforms to Levelt’s propositions

To further test the psychophysical validity of our neurobiologically detailed model, we simulated the experimental conditions described by Levelt’s modified propositions (Brascamp et al., 2015) – a set of four statements that compactly summarise the relationship between stimulus strength (e.g., luminance contrast, colour contrast, stimulus motion) and the average dominance duration of each stimulus percept. Here, we focus on the modified second and fourth propositions, as they constitute the “core laws” of rivalry and incorporate recent psychophysical findings (propositions one and three are consequences of proposition two; Brascamp et al., 2015).

Levelt’s modified second proposition states that increasing the difference in stimulus strength between the two eyes will principally increase the average dominance duration of the percept associated with the stronger stimulus (Brascamp et al., 2015; Leopold & Logothetis, 1996; Logothetis & Leopold, 1996). To simulate Levelt’s second proposition, we decreased the spike rate entering one side of the ring from 1350 to 1200 [Hz] in steps of 50 [Hz] across simulations. In line with predictions (Fig 3F), the average dominance duration of the percept corresponding to the stronger stimulus showed a steep increase from ∼3.25 [s] with matched input, to ∼ 6 [s] with maximally different input whilst the average dominance duration on the side of the weakened stimulus decreased comparatively gradually to ∼1.9 [s] with maximally different inputs.

According to Levelt’s modified fourth proposition, increasing the strength of the stimulus delivered to both eyes will increase the average perceptual reversal rate (i.e., decrease dominance durations; Brascamp et al., 2015), a finding that has been replicated across a wide array of experimental settings (Bonneh et al., 2014; Brascamp et al., 2006; Buckthought et al., 2008; Meng & Tong, 2004). To simulate Levelt’s fourth proposition, we ran a series of simulations in which we increased the spike rate of the external drive in steps from 1300 to 1500 [Hz] in steps of 50 [Hz] across simulations. Again in line with predictions (Fig 3G), the perceptual alternation rate increased with input strength, starting at ∼7.75 alternations per minute at the second weakest stimulus strength (1350 [Hz]) and increasing to ∼15 alternations per minute for the strongest stimulus (1500 [Hz]). Interestingly, along with a number of meanfield models of rivalry (Shpiro et al., 2007), our model predicts a deviation from Levelt’s fourth proposition for very low stimulus values with an uptick in alternation rate occurring at the lowest external dive value (1300 [Hz]). Encouragingly, there is some initial evidence that deviations from Levelt’s fourth law may be present in human psychophysical data (Brascamp et al., 2015).

To help ensure that the simulation results were not biased by finite size effects or other simplifying assumptions such as the all-to-all connectivity of the cortical ring, or the 50/50 excitatory/inhibitory neuron ratio, we show in supplementary material S5 that a scaled-up version model consisting of 2000 cortical neurons with sparse connectivity, and an 80/20 excitatory/inhibitory neuron ratio (i.e. consistent with Dale’s law), also produces a Gamma distribution of dominance durations, and is consistent with Levelt’s second and fourth propositions.

We thus confirmed that our neurobiologically detailed model of the matrix thalamus - L5PT loop is capable of reproducing Levelt’s propositions, which together with the right-skewed distribution of dominance durations, show the consistency of our model with the psychophysical “laws” known to govern visual rivalry.

### Generating testable predictions through in silico electrophysiology

Binocular rivalry is thought to depend in part on the substantial degree of binocular overlap in humans (∼120°), however the lateral position of the eyes in mice leaves only ∼40 ° of binocular overlap (Poort & Meyer, 2021). For this reason, there are no current mouse models of binocular rivalry, however there are monocular variants of visual rivalry, namely plaid perception, that can be studied the mouse model (Bogatova et al., 2024; Palagina et al., 2017). Crucially, plaid perception, like binocular rivalry, conforms to Levelt’s laws (Brascamp et al., 2015; Hupé et al., 2019) and also has a right skewed distribution of dominance durations that is well fit by a Gamma distribution (Bogatova et al., 2024). We hypothesise, therefore, that the principles underlying the simulation of binocular rivalry in our model will also describe other forms of visual rivalry (such as plaid perception), offering a plausible means to test cellular level predictions derived through simulation. To this end, we next ran a series of perturbation experiments, with the aim of interrogating the novel burst-dependent mechanism of perceptual dominance by mimicking the optogenetic and pharmacological experiments carried out in threshold-detection studies (Takahashi et al, 2016; 2020), in the context visual rivalry. As perceptual dominance depends on the formation and maintenance of a burst-dependent persistent state, we hypothesised that artificially exciting the apical dendrites would result in an increase in the average dominance duration of the excited population, and artificially inhibiting the apical dendrites and thalamus would result in a decrease in the dominance durations for the inhibited populations.

Due to the fact that the distance to bifurcation of the dominant population fluctuates around zero, we predicted that exciting the apical dendrites would increase the proportion of the population above the bifurcation at B1, thereby reducing the probability with which a fluctuation in somatic drive would lead to a sizable enough drop in the proportion of the population below B1 to release the competing population from inhibition. This should, therefore, result in an increase in the frequency of long dominance duration events, thereby increasing the mean and the spread of the distribution. Equivalently, we predicted that inhibiting the apical dendrites would reduce the proportion of the population above B1, making it more likely that transient fluctuations in somatic drive would allow the competing population to escape from inhibition, reducing the occurrence of long dominance duration events, thereby reducing both the mean and the spread of the distribution of dominance durations. Finally, based on the results of the threshold detection simulations we predicted that thalamic inhibition would have an analogous effect on dominance durations to apical dendrite inhibition but would be mediated by a reduction in the coupling probability. To test these hypotheses, we conducted two *in silico* experiments analogous to the conditions described by Levelt’s modified propositions but instead of manipulating the external drive entering the somatic compartment of L5PT cells we manipulated the amplitude of simulated causal perturbations to the L5PT apical compartment and thalamus (Fig 4A & 5A).

**Fig 4.**
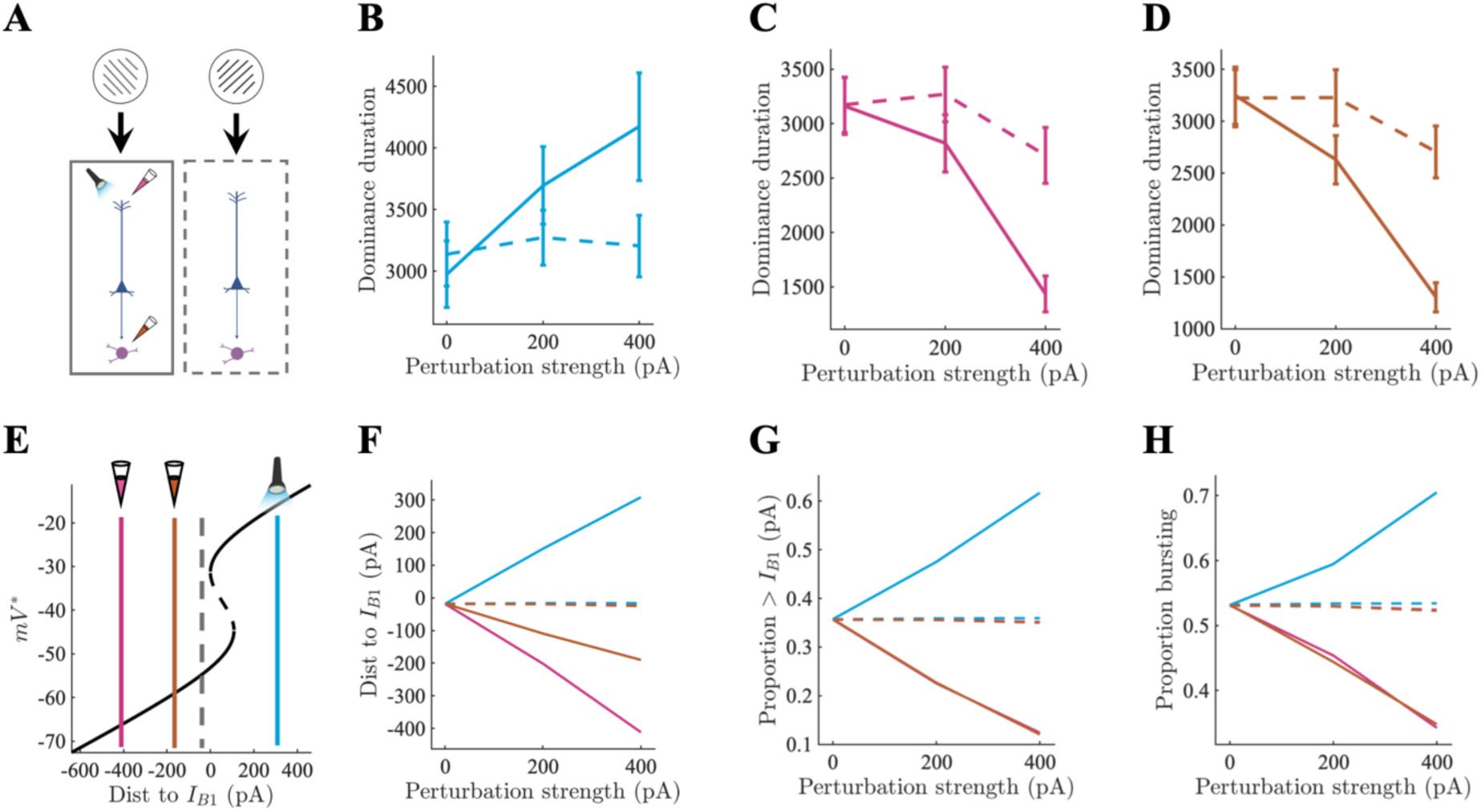
*In silico* electrophysiology reveals degenerate mechanisms of perceptual dominance (asymmetric perturbations). A) Causal perturbations to one half of the thalamocortical circuit underlying visual rivalry consisting of optogenetic excitation of the apical compartment (blue), pharmacological inhibition of the apical compartment (pink), and pharmacological inhibition of the thalamus (orange). B-D) Average dominance duration of perturbed (solid), and unperturbed (dashed) populations. Error bars show SEM. E) Average distance to bifurcation point at B_1_ shown on bifurcation diagram for perturbed (solid) and unperturbed (dashed) populations during periods of perceptual dominance with 400 [pA] perturbation strength. F) Average distance to bifurcation point at B_1_ for perturbed (solid) and unperturbed (dashed) populations during periods of perceptual dominance. G) Proportion of population above bifurcation point at B_1_ during periods of perceptual dominance. H) Proportion of population in bursting regime during periods of perceptual dominance.

In the first set of experiments, we simulated optogenetic excitation and pharmacological inhibition of one of the two competing populations by adding a constant current (± 200, 400 [pA]) to all of the target variables (i.e. apical compartment or thalamic neurons) on one side of the ring (Fig 4A). In line with predictions, we found that the average dominance duration of the excited population (Fig 4B) increased, the distance to bifurcation decreased (Fig 4E-F), the proportion of the population above the critical point at B1 increased (Fig 4G), and the proportion of the population in the bursting regime increased (Fig 4H). The dominance durations and neuronal dynamics of the unexcited population remained relatively unchanged.

Similarly, inhibition of both the apical dendrites and thalamus reduced the average dominance duration of the inhibited population whilst the uninhibited population was again relatively unchanged (Fig 4C-D). As in the threshold detection simulations, inhibition of the apical dendrites led to a large increase in the distance to B1 compared to thalamic inhibition (Fig 4E-F) which primarily affected the inter-compartment coupling probability (supplementary material S6A). Both apical dendrite and thalamic inhibition led to almost identical reductions in the proportion of the population above the critical point at B1 (Fig 4G), and the proportion of the population in the bursting regime (Fig 4H). The uninhibited population again remained relatively constant across all of the neuronal measures (the small drop in dominance durations of the uninhibited population for the 400 [pA] inhibitory perturbations is due to the adaptation variable having less time to recover). As predicted the spread of the distribution of dominance durations increased with the amplitude of excitatory perturbation and decreased under inhibition (supplementary material S6B).

In the second set of experiments, we simulated optogenetic excitation and pharmacological inhibition of both competing neuronal populations simultaneously by adding a constant current (± 200, 400 [pA]) to all of the target variables on the ring (Fig 5A). Again, in line with predictions, the speed of rivalry (i.e., the number of perceptual alternations per minute of simulation time) decreased as a function of apical dendrite excitation (Fig 5B). Excitation also decreased the average distance to B1 (Fig 5E-F), increased the proportion of the population above the critical point at B1 (Fig 5G), and increased the proportion of the population in the bursting regime (Fig 5H).

**Fig 5.**
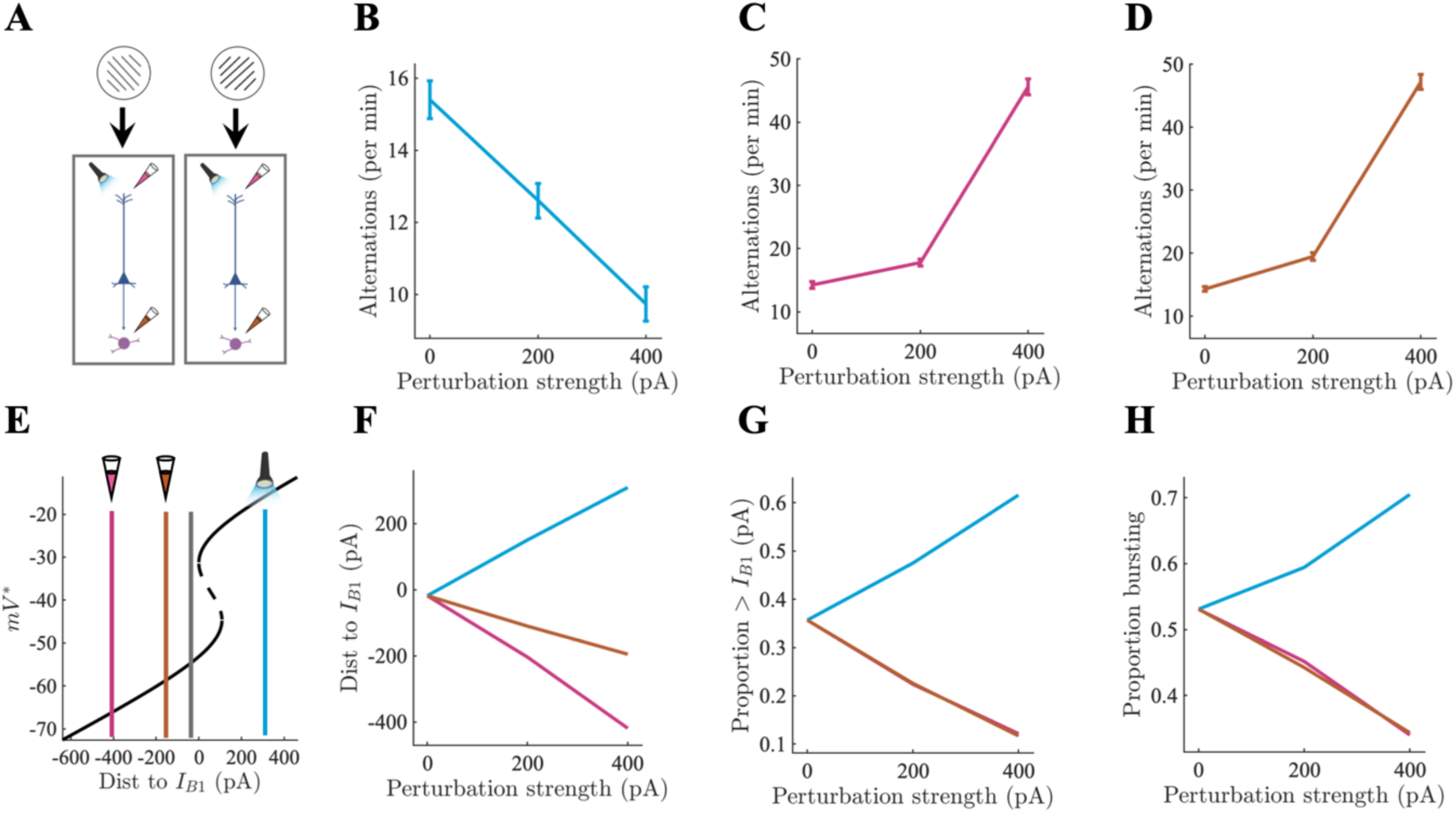
*In silico* electrophysiology reveals degenerate mechanisms of perceptual dominance (symmetric perturbations). A) Causal perturbations to full thalamocortical circuit underlying visual rivalry. Colours same as above. B-D) Perceptual alternations per minute of simulation time across perturbation types. Error bars show SEM. E) Average distance to bifurcation point at B_1_ shown on bifurcation diagram for perturbed (blue, orange, pink) and unperturbed (grey) simulations during periods of perceptual dominance at 400 [pA]. F) Average distance to bifurcation point at B_1_ for full network perturbations during periods of perceptual dominance. G) Average proportion of population above bifurcation point at B_1_ for full network perturbations during periods of perceptual dominance. H) Average proportion of population in bursting regime for full network perturbations during periods of perceptual dominance.

In contrast, inhibition of the apical dendrites and thalamus both increased the speed of rivalry (Fig 5C-D). Inhibition of the apical dendrites led to a large increase in the distance to the critical point at B1 compared to thalamic inhibition (Fig 5E-F), but both apical dendrite inhibition and thalamic inhibition reduced the proportion of the population above the critical point at B1 (Fig 5G), and the proportion of the population in a bursting regime (Fig 5H) through reductions in the thalamus mediated inter-compartment coupling probability (supplementary material S6C). As with the asymmetric perturbation simulations, inhibition exerted a much larger effect on the speed of rivalry than excitation. Finally, again in line with our predictions, the spread of the distribution of dominance durations increased under excitatory perturbation of the apical dendrites and decreased under inhibition of the apical dendrites and thalamus (supplementary material S6D).

Together, these *in silico* electrophysiological experiments provide important testable (and explainable) hypotheses for future experiments that although not testable in any existing data sets are well within the purview of modern systems neuroscience providing an opportunity to conduct precise theory-driven tests of the model.

## Discussion

The study of perceptual awareness in human participants and animal models has so far proceeded largely in parallel – the former exploring the largescale neural dynamics and behavioural signatures of perceptual awareness across a rich array of experimental settings, and the latter characterising the cellular circuitry of perception in exquisite detail, and with precise causal control, but with only limited links to higher level perceptual phenomena (He, 2023). Leveraging a neurobiologically detailed model of the matrix thalamus – L5PT loop, we have shown that a potential circuit-level mechanism of tactile perceptual awareness discovered in a mouse model of tactile awareness (Aru et al., 2019, 2019; Takahashi et al., 2016, 2020) generalises to visual rivalry, thus providing a roadmap for the linking circuit level mechanisms studied in animal models to the behavioural signatures of perceptual awareness studied in human participants.

The balance of neurobiological detail and interpretability offered by our model allowed us to reproduce the threshold-detection results of Takahashi et al (2016, 2020) and interrogate the mechanisms underlying the experiments in a manner that would be impossible *in vivo*. In particular, examination of the model’s dynamics under simulated causal perturbations to the circuit revealed a degenerate dynamical mechanism for controlling the threshold for perceptual awareness. Excitation of the apical compartment reduced the distance to bifurcation in the apical compartment, thus increasing the probability that each cell could generate a Ca^2+^ plateau potential switching the soma into a bursting regime. This resulted in an increase in the baseline and stimulus-evoked spike count, and correspondingly, led to a reduction in the model’s perceptual threshold. Inhibition of the apical compartment and thalamus resulted in comparable downward shifts in the baseline and stimulus-evoked spike count, leading to increases in the model’s perceptual threshold. Importantly, however, the neural mechanisms underlying the increases in perceptual threshold were distinct: inhibiting the apical compartment increased the distance to bifurcation, thus reducing the probability with which each cell would generate a Ca^2+^ plateau potential, whereas inhibiting the thalamus reduced the inter-compartment coupling. Both mechanisms, however, led to comparable reductions in the proportion of cells in the bursting regime explaining the comparable increase in perceptual thresholds, suggesting that it is the emergent action of the corticothalamic circuit as a whole, rather than single cells within the circuit, that are responsible for perceptual awareness.

The degenerate mechanisms underlying the threshold for perceptual awareness combined with the operational definition of perceptual awareness in the threshold detection task (in terms of psychometric functions) points to a conceptually important point about the role of bursts in the model, and potentially, the empirical data itself. Specifically, controlling the ease with which a cell can burst through optogenetic and pharmacological perturbation is simply a means for controlling how easily a stimulus can evoke reverberant activity in corticocortical and thalamocortical loops which, in the simple case of threshold detection, constrains the extent to which stimulus evoked activity can stand out against a background of noise driven fluctuations.

We next showed that the same thalamus-gated burst-dependent mechanism underlying perceptual awareness in simulations of the tactile threshold detection task also determines perceptual dominance in simulations of visual rivalry. Specifically, perceptual dominance is initiated by a succession of regular spikes and maintained through the formation of a transiently stable burst-dependent persistent state characterised by reliable coupling between apical and somatic compartments. This allows the apical compartment to generate temporally extended plateau potentials in a large subset of the dominant population reliably switching the L5PT soma from a regular spiking to a bursting regime. Perceptual dominance is then maintained until the slow hyperpolarising adaptation current accumulates to a sufficiently high level that the dominant population is no longer able to maintain inhibit the competing and a perceptual switch ensues.

Importantly, the model conforms to Levelt’s modified propositions. Originally proposed in 1965 (Levelt, 1965), “Levelt’s laws” have proven to be remarkably robust needing only minor modification and contextualisation (Brascamp et al., 2015) and have, therefore, served as a benchmark for computational models of visual rivalry (e.g., Grossberg et al., 2008; Laing & Chow, 2002; Shpiro et al., 2007; Wilson, 2007). Together with the right-skewed (Gamma) distribution of dominance durations the consistency of our model with Levelt’s propositions provides an *in silico* conformation of the hypothesis that pulvinar – L5PT loops in visual cortex may play an analogous role to POm – L5PT loops in barrel cortex. This is a minimal but necessary first step in testing the hypothesis that reverberant activity in matrix thalamus – L5PT loops is a necessary component part in a domain general mechanism of perceptual awareness.

Having validated our model against psychophysical benchmarks, we next sought to interrogate the novel thalamus-gated burst-dependent mechanism of perceptual dominance by emulating the optogenetic and pharmacological experiments carried out by Takahashi et al (2016, 2020) in the context of visual rivalry. Under conditions of visual rivalry, the simulated causal perturbations are similar to the conditions described by Levelt’s propositions, but instead of manipulating the strength of the external stimulus we manipulated the strength apical compartment excitation/inhibition, or thalamic inhibition, highlighting the unique contribution of these neurobiological components to visual rivalry. Across asymmetric and symmetric perturbations excitation of the apical compartment slowed perceptual alternations (i.e., increased dominance durations) by increasing the proportion of the population able to sustain temporally extended Ca^2+^ plateau potentials and remain in a transiently-stable bursting regime, whereas inhibition of both the apical dendrites and thalamus had the opposite effect. Although technically difficult, these simulated experimental manipulations are well within the purview of modern experimental techniques and therefore represent a means of causally testing the predictions of our model. Importantly, the simulation of these experimental perturbations would not be possible in any existing models of rivalry, even those at the spiking level (e.g. Laing & Chow, 2002; Wang et al., 2020; Wilson, 2003), as they focus on the minimal conditions for rivalry in point-neuron models of cortical interaction. The inclusion of a dual compartment model of L5PT cells, and an explicit thalamic population, was, therefore, required in order to make contact with the results of Takahashi et al (2016, 2020).

In addition to the predicted effect of causal perturbations on visual rivalry, our model generates a number of more straightforward correlational predictions. Specifically, matrix-rich higher-order thalamic nuclei with recurrent connections to sensory cortex, such as the pulvinar, should be selective for perceptual awareness rather than physical stimulation, a prediction supported by both human neuroimaging (Qian et al., 2023; Seo et al., 2022) and non-human primate electrophysiology (Wilke et al., 2009). Similarly, synchronous bursting activity in deep layers of cortex, specifically layer 5b which contain the soma of ttL5PT cells, should likewise be selective for perceptual awareness rather than physical stimulation a prediction that, with the advent of primate Neuropixels (Trautmann et al., 2023), is also readily testable. Finally, in the context of visual rivalry, perceptual dominance should be characterised by elongated Ca^2+^ plateau potentials in the apical dendrites of L5PT cells (located in L1) in cells selective for the dominant percept, a prediction testable in mouse models of visual rivalry (e.g. Bogatova et al., 2024).

We anticipate that the cellular conditions for awareness explored in this paper are likely to have consequences for the largescale correlates of awareness. Indeed, we venture that at the level of large scale brain networks diffuse matrix-thalamus gated bursting may play a key role in the formation of a quasi-critical regime (Müller et al., 2020, 2023) allowing single nodes in a network to transiently escape from a tight E/I balanced state. This permits stimulus information to rapidly propagate across the cortical sheet whilst also maintaining stability at the level of the whole network (Müller et al., 2020) effectively modulating the gain of interareal connectivity in line with previous computational models of pulvinar-cortical interactions in cognitive tasks (Jaramillo et al., 2019). Indeed, efforts to test the largescale consequences of the cellular level mechanisms interrogated in this paper are already very much underway. Biophysical modelling of source-localised MEG data showed that auditory awareness evoked activity was best fit by increased input to superficial layers of the cortical column consistent with the projections of matrix-type higher-order thalamus (Pujol et al., 2023).

As has been noted elsewhere (c.f. Aru et al., 2020; Storm et al., 2024), the circuit level conditions for awareness explored here fit well with many of the major neuronal theories of consciousness. The diffuse projections of the matrix-type thalamus may be a circuit level mechanism underlying the non-linear and widespread “ignition” response proposed by global neuronal workspace theory to underlie the transition from unconscious to conscious processing (Benitez et al., 2023; Cortes et al., 2023; Klatzmann et al., 2022; Mashour et al., 2020). The improvement in signal-to-noise ratio associated with bursting aligns with signal detection theoretic versions of higher-order theory (Lau, 2007), and the recurrent nature of matrix thalamus – L5PT loops could be considered a thalamocortical extension of the currently corticocentric recurrent processing theory (Lamme, 2006). In addition, in previous work we have shown that diffuse matrix-type control of bursting in a sheet of L5PT cells maximises an approximate measure of integrated information (Munn, Müller, Aru, et al., 2023), in line with integrated information theory (Albantakis et al., 2023). We speculate that exploring the interaction between the cellular conditions for awareness interrogated in this paper, and the topology of largescale brain networks, may be of crucial importance in resolving the ongoing debate between the theories described above regarding the macroscale network conditions necessary for awareness.

To strike the right balance between neurobiological detail and interpretability, we made a number of simplifying assumptions that place some limitations on our model. Most notably, we did not include L2/3 pyramidal neurons – which are arguably the primary source of long distance horizontal connections in the cortex (Douglas & Martin, 2004) and arguably cross column inhibition (Qian et al., 2023) – nor a core thalamic population which forms a targeted recurrent loops with L4 and L6 of cortex (Harris & Shepherd, 2015) preventing us from performing systematic perturbation experiments on our model highlighting the precise function of L5PT cells and higher-order matrix thalamus in a more realistic cortical microcircuit. We also did not include time delays between our corticocortical or thalamocortical connections preventing our model from providing a realistic model (e.g., Tahvili & Destexhe, 2023) of time-frequency components of common electrophysiological measures such as local field potentials. Finally, our model has only a single hierarchical level preventing us from making contact with evidence showing a potential prefrontal contribution to perceptual switches (Dwarakanath et al., 2020; Kapoor et al., 2020).

Our model is, of course, only a first step towards a formal characterisation of the minimal neurobiological mechanisms underlying perceptual awareness. Extending the model, and modelling strategy more generally, to new paradigms such as backward masking (Gale et al., 2024), will be of paramount importance in the progression of the field as mouse models and the tools of systems neuroscience are brought into contact with the sophisticated psychophysical paradigms used to study the behavioural signatures of awareness in humans.

## Materials and Methods

### Thalamocortical spiking neural network

The neuronal backbone of the model consists of a (novel) dual compartment model of L5PT neurons, fast spiking interneurons (basket cells), and thalamic cells. The dynamics of basket cells, thalamic cells, and the somatic compartment of L5PT cells (Fig 1A-C) were described by Izhikevich quadratic adaptive integrate and fire neurons, a hybrid dynamical system that is capable of reproducing a wide variety spiking behaviour whilst still being highly efficient to integrate numerically (Izhikevich, 2003, 2004, 2006). The Izhikevich neuron consists of the following two-dimensional system of ODEs:

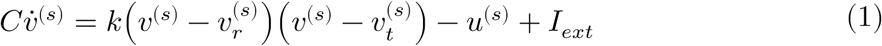

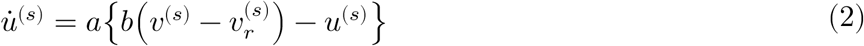

with reset conditions: if 𝑣 ≥ 𝑣_𝑝𝑒𝑎𝑘_ then 𝑣 → 𝑐, 𝑢 → 𝑢 + 𝑑. The equations are in dimensional form giving the membrane potential (including the resting potential 𝑣_𝑟_, spike threshold 𝑣_𝑡_, and spike peak 𝑣_𝑝𝑒𝑎𝑘_, and reset c), input 𝐼_𝑒𝑥𝑡_, time 𝑡, and capacitance 𝐶, biophysically interpretable units (mV, pA, mS, and pF respectively). The remaining four parameters 𝑘, 𝑎, 𝑏, and d, are dimensionless and control the sharpness of the quadratic-nonlinearity, the timescale of spike adaptation, the sensitivity of spike adaptation to sub-threshold oscillations, and the magnitude of the spike reset adaptation variable. Crucially, Izhikevich (Izhikevich, 2006; Izhikevich & Edelman, 2008) fit parameters for a large class of cortical and sub-cortical neurons, thus affording our model a high degree of neurobiological plausibility while greatly reducing the number of free parameters.

The apical compartment of the L5PT neuron consists of a two dimensional non-linear system introduced by Naud and colleagues (Naud & Sprekeler, 2018) as a phenomenological model of the Ca^2+^ plateau potential in the apical dendrites of L5PT neurons.

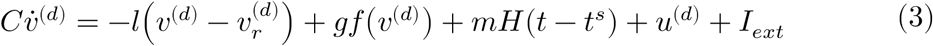

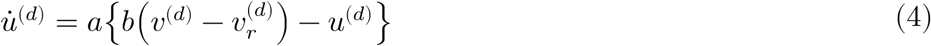

Where 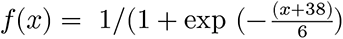 describes the regenerative non-linearity underlying the Ca^2+^ plateau potential, and 𝐻(𝑡 − 𝑡^𝑠^) denotes a square wave function of unitary amplitude describing the backpropagating action potential (delayed by 0.5 [ms] and lasts for 2 [ms]) with 𝑡^𝑠^ denoting the somatic compartment spike time. The parameters 𝑙, 𝑔, 𝑚, denote the leak conductance [nS], amplitude of regenerative non- linearity [pA], and amplitude of the back propagating action potentials [pA] respectively. The model and parameters were derived from a more complex model of Ca^2+^ spikes built to predict *in vitro* L5PT spike times (Naud et al., 2014).

To simulate key observations from empirical experiments, we coupled the compartments together so that sodium spikes in the somatic compartment triggered a back propagating action potential affecting the apical compartment through the square wave function 𝐻(𝑡 − 𝑡^𝑠^). In turn, plateau potentials in the apical compartment controlled the reset conditions of the somatic compartment. We leveraged the insight (Izhikevich, 2003; Munn, Müller, Aru, et al., 2023; Munn, Müller, Medel, et al., 2023) that the difference between regular spiking and intrinsic bursting can be modelled by changing the reset conditions of equations (1) and (2), raising the reset voltage (increasing 𝑐) taking the neuron closer to threshold, and reducing the magnitude of spike adaptation (decreasing 𝑑). Whenever the membrane potential in the apical compartment exceeded −30 mv the reset conditions changed from regular spiking to bursting parameters. This allowed us to reproduce the transient change in dynamical regime in L5PT cells that occurs when they receive coincident apical and basal drive. Parameters values for each neuron/compartment are given in table 1.

**Table 1.**
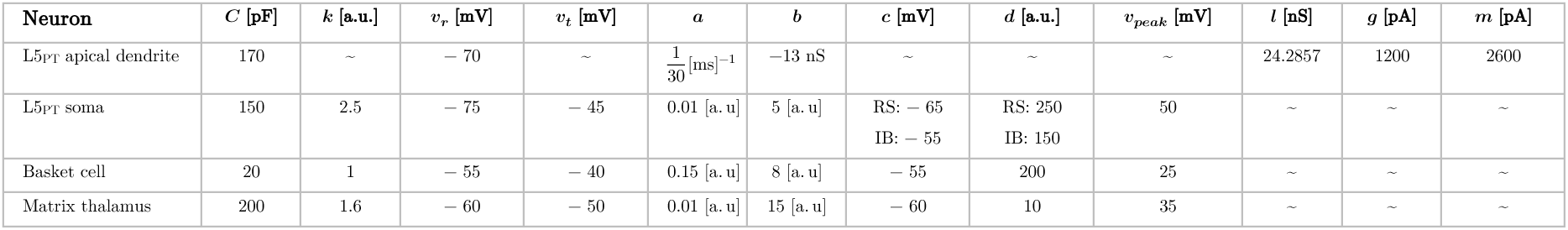
Parameters for each neuron L5_PT_ apical dendrite parameters were taken from Naud and Sprekeler, (2018). L5_PT_ soma parameters were modified from the model of intrinsic bursting (p.290) described in Izhikevich (2006). Basket cell (fast spiking interneuron), and matrix thalamus parameters were taken from Izhikevich and Edelman (2008).

Based on the finding that communication between apical dendrites and the soma of L5PT cells requires depolarising input from the matrix thalamus to the “apical coupling zone” in L5a (Suzuki & Larkum, 2020) we made back propagating action potentials and Ca^2+^ driven parameter switches depend stochastically upon a phenomenological model of excitatory dynamics in the apical coupling zone described by the saturating linear system shown in equation (5).

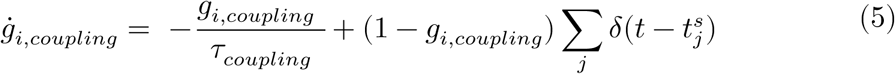

Coupling was driven by thalamic spikes (where 𝑡^𝑠^ denotes the time that the thalamic neuron passes the threshold 𝑣 ≥ 𝑣_𝑝𝑒𝑎𝑘_) and the decay constant 𝜏_𝑐𝑜𝑢𝑝𝑙𝑖𝑛𝑔_ was taken from work estimating the decay of the post synaptic excitatory effects of metabotropic glutamate receptors (Greget et al., 2011) which have been shown empirically to mediate inter-compartmental coupling in L5PT cells (Suzuki & Larkum, 2020). By design, the dynamics of the coupling variable varied between 0 and 1 and governed the probability with which back propagating action potentials would reach the apical compartment and the probability with which a Ca^2+^ spike would lead to a switch in the soma reset parameters.

Based on previous spiking neural network models of rivalry (Laing et al., 2010; Laing & Chow, 2002; Wang et al., 2020) the cortical component of the network had a one-dimensional ring architecture. Each point on the ring represents an orientation preference with one full rotation around the ring corresponding to a 180^°^ visual rotation. This mirrors the fact that a 180^°^ rotation of a grating results in a visually identical stimulus and also ensures periodic boundary conditions. The cortical ring contained 90 L5PT neurons and 90 fast spiking interneurons. Each pair of excitatory and inhibitory neurons was assigned to an equidistant location on the ring (unit circle) giving each neuron a 2^°^ difference in orientation preference relative to each of its neighbours. The (dimensionless) synaptic weights 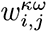 connecting neurons (𝐸 → 𝐸, 𝐼 → 𝐸, and 𝐸 → 𝐼), were all-to-all with amplitude decaying as a function of the Euclidean distance 𝑑_𝑖,𝑗_ between neurons (equations (6)) according to a spatial Gaussian footprint (equation (7)).

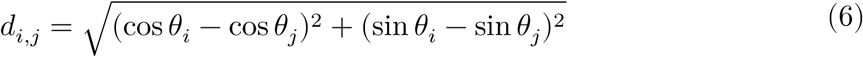

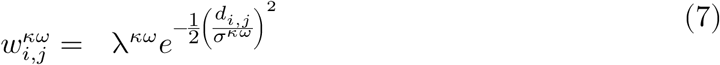

Where 𝜃 is the location of the neuron on the unit circle, 𝜅 and 𝜔 denote the pre- and post-synaptic neuron type (i.e. 𝐸 → 𝐸), λ controls the magnitude of the synaptic weights, and 𝜎^𝜅𝜔^ the spatial spread. In line with empirical constraints inhibitory coupling had a larger spatial spread than excitatory to excitatory coupling (Naka & Adesnik, 2016). Each thalamic neuron received input from 9 cortical neurons and then projected back up to the apical dendrites of the same 9 cortical neurons recapitulating the diffuse projections of higher-order thalamus onto the apical dendrites of L5PT neurons in layer 1 (Mease & Gonzalez, 2021; Shepherd & Yamawaki, 2021). For simplicity we set projections to and from the thalamus to a constant value (e.g., 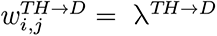). For the sake of computational efficiency we also neglected differences in rise time between receptor types which allowed us to model receptor dynamics with a first-order linear differential equation (equation 8) with decay (𝜏_𝑑𝑒𝑐𝑎𝑦_) constants chosen to recapitulate the dynamics of inhibitory (GABAA), and excitatory (AMPA and NMDA) synapses (Dayan & Abbott, 2005; Gerstner et al., 2014).

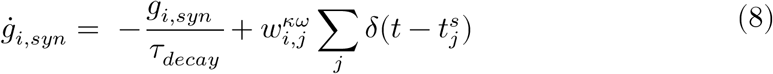

Where, as above, denotes 𝑡^𝑠^ the time that the neuron passes the threshold 𝑣 ≥ 𝑣_𝑝𝑒𝑎𝑘_. The conductance term entered into the input 𝐼_𝑒𝑥𝑡_ through the relation 𝐼_𝑖,𝑠𝑦𝑛_ = 𝑔_∞_𝑔_𝑖_(𝑡)_𝑠𝑦𝑛_(𝐸_𝑠𝑦𝑛_ − 𝑣_𝑖_(𝑡)) where 𝐸_𝑠𝑦𝑛_ is the reverse potential of the synapse. Following Izhikevich and Edelman (2008) we set 𝑔_∞_ to 1 for GABAA and AMPA synapses, and 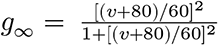 for NMDA synapses. To prevent artificial distortions of the spike shape that can occur during parameter sweeps that push the model outside its normal operating regime we clipped individual NMDA conductances to a maximum value of 85 [nS].

For the threshold detection simulations, the somatic compartment of each L5PT cell received 600 [Hz] of independent (Poisson) external drive and apical compartments received 50 [Hz] of external drive. The whisker deflection was simulated by a pulse of constant amplitude varying between 0 – 350 [pA] lasting 200 [ms] and weighted by the spatial Gaussian shown in equation (9) where N is the neuron at the centre of the pulse and 𝜎^𝑇𝐷^ the spatial spread.

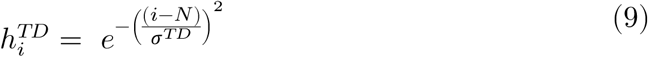

For the visual rivalry simulations, separate monocular inputs targeting the somatic compartment of L5PT cells were modelled with two independent Poisson processes (representing input from the left and right eyes in the case of binocular rivalry or left and right movement selective populations in the case of plaid perception) with rates varying between 1200 and 1800 [Hz] depending on the simulation. The external drive was weighted by the spatial Gaussian shown in equation (10) centred on neurons 90^°^ apart on the ring abstractly corresponding to the orthogonal grating stimuli commonly employed in binocular rivalry experiments.

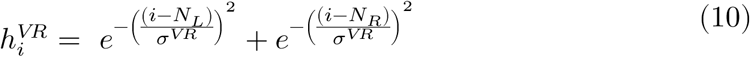

Here 𝑖 denotes the index of the 𝑖th neuron, 𝑁_𝐿_ and 𝑁_𝑅_ control the orientation of the stimulus delivered to the left and right eyes, and 𝜎^𝑉𝑅^ the spatial spread.

To capture the slow hyperpolarising current that traditionally governs switching dynamics in models of bistable perception (Wilson & Cowan, 2021), the somatic compartment of each L5PT cell was coupled to a phenomenological model of slow hyperpolarising Ca^2+^ mediated K^+^ currents (McCormick & Williamson, 1989) which entered into the external drive term for each cell (i.e. 𝐼_𝑖,𝑎𝑑𝑎𝑝𝑡_ = 𝑔_𝑖_(𝑡)_𝑎𝑑𝑎𝑝𝑡_(𝐸_𝑎𝑑𝑎𝑝𝑡_ − 𝑣_𝑖_(𝑡))) with dynamics given by equation (10).

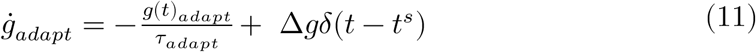

Where Δ𝑔 denotes the contribution of each spike to the hyperpolarising current, and 𝜏_𝑎𝑑𝑎𝑝𝑡_ the decay constant.

Rather than fit the parameters of our model to individual experimental findings, which permits substantial degrees of freedom and risks overinterpretation of idiosyncratic aspects of individual experiments, we instead elected to challenge a single model to qualitatively reproduce a wide array of experimental findings with a minimal set of parameter changes carefully chosen to reflect experimental manipulations and perturbations. Specifically, we initialised the connectivity parameters such that: 1) when the model received a background drive the conductances were approximately E/I balanced with a coefficient of variation > 1, corresponding to an asynchronous irregular regime (Destexhe, 2009); 2) inhibitory connections on the cortical ring had broader (Gaussian) connectivity than excitatory connections generating a winner-take-all regime when the model received two “competing” inputs to opposite sides of the cortical ring; and 3) a slow hyperpolarising current was added to the somatic compartment of each L5PT cell destabilizing the winner-take-all attractor states leading to spontaneous switches between transiently stable persistent states with an average duration in the experimentally observed range for binocular rivalry. Parameters for the model components described by equations (5) – (11) are supplied in table 2.

**Table 2.** Parameter description, values and units for the model components described by equations (5- 11).

| Parameter | Description | Value | Units |
| --- | --- | --- | --- |
| $\lambda_{AMPA}^{E \rightarrow E}$ | Amplitude of excitatory to excitatory coupling for (AMPA) | $\frac{6.125}{\sigma^{E \rightarrow E} \sqrt{2\pi}}$ | a.u. |
| $\lambda_{NMDA}^{E \rightarrow E}$ | Amplitude of excitatory to excitatory coupling (NMDA) | $\frac{1.225}{\sigma^{E \rightarrow E} \sqrt{2\pi}}$ | a.u. |
| $\lambda^{E \rightarrow I}$ | Amplitude of excitatory to inhibitory coupling (NMDA and AMPA) | $\frac{1}{\sigma^{E \rightarrow I} \sqrt{2\pi}}$ | a.u. |
| $\lambda^{I \rightarrow E}$ | Amplitude of inhibitory to excitatory coupling (GABA<sub>A</sub>) | $\frac{5}{\sigma^{I \rightarrow E} \sqrt{2\pi}}$ | a.u. |
| $\lambda^{E \rightarrow TH}$ | Constant excitatory to thalamic coupling constant (AMPA only) | 4 | a.u. |
| $\lambda_{AMPA}^{TH \rightarrow D}$ | Constant thalamic to apical dendrite coupling (AMPA) | 10 | a.u. |
| $\lambda_{NMDA}^{TH \rightarrow D}$ | Constant thalamic to apical dendrite coupling (NMDA) | 10 | a.u. |
| $\sigma^{E \rightarrow E}$ | Spread of excitatory to excitatory coupling | 0.5 | a.u. |
| $\sigma^{E \rightarrow I}$ | Spread of excitatory to inhibitory coupling | 2 | a.u. |
| $\sigma^{I \rightarrow E}$ | Spread of inhibitory to excitatory coupling | 2 | a.u. |
| $\tau_{decay}$: AMPA | Decay time of AMPA conductance | 6 | ms |
| $\tau_{decay}$: GABA<sub>A</sub> | Decay time of GABA<sub>A</sub> conductance | 6 | ms |
| $\tau_{decay}$: NMDA | Decay time of NMDA conductance | 100 | ms |
| $\tau_{coupling}$ | Decay time of apical coupling zone | 800 | ms |
| $\tau_{adapt}$ | Decay time of adaptation current | 2000 | ms |
| $\Delta g$ | Contribution of each spike to adaptation current | 0.065 | nS |
| $\sigma^{TD}$ | Spatial spread of external drive | 20 | a.u. |
| $\sigma^{VR}$ | Spatial spread of apical drive | 18 | a.u. |
| $E_{Excitatory}$ | Reverse potential of excitatory synapses | 0 | mV |
| $E_{Inhibitory}$ | Reverse potential of inhibitory synapses | -75 | mV |
| $E_{Adapt}$ | Reverse potential of adaptation currents. | -80 | mV |

The equations were integrated numerically in MATLAB 2023b. The apical compartment was integrated with a standard forward Euler scheme. All other compartments were integrated using the hybrid scheme for conductance based models introduced by Izhikevich (2010). All simulations used a step size of 0.1 [ms] and were run for 30 [s]. Unless stated otherwise, all simulation results were averaged over a minimum of 30 random seeds.

### Distance to bifurcation

To obtain a closed form expression for the distance to bifurcation in the apical compartment (equations 3-4), we leveraged the fact that saddle node bifurcations occur when the nullclines (𝑣̇^(𝑑)^ = 𝑢̇^(𝑑)^ = 0) of the system intersect tangentially (Strogatz, 2018). That is, the nullclines and derivative of the nullclines must be equal leading to the following two requirements (where we have absorbed the term describing back propagating action potentials 𝐻(𝑡 − 𝑡^𝑠^) into the external drive 𝐼_𝑒𝑥𝑡_ which we treat as a constant).

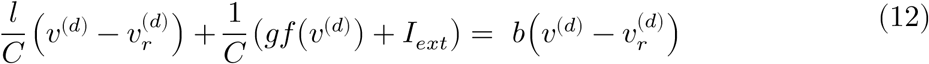

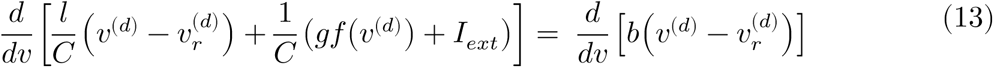

We used equation (13) to solve for 𝑣^(𝑑)^ giving 𝑣^(𝑑)∗^ = [−44.6601, −31.3399]. We then substituted 𝑣^(𝑑)∗^ back into equation (12) to solve for 𝐼_𝑒𝑥𝑡_ yielding the value of the external current at each of the two bifurcations IB1 = 538.911 [pA] and IB2 = 647.375 [pA] (corresponding to points at which the linear adaptation current nullcline intersects tangentially with the left and right knees of the cubic membrane potential nullcline see supplementary material S1B-D).

### Psychometric and neurometric functions

To obtain a measure of response probability from our model comparable to the psychometric functions in Takahashi et al (2016, 2020) we took a two pronged approach. First, for all stimulus intensities including stimulus absent trials (when the model only received a background drive) we calculated the frequency with which the spike count in the 1000ms post stimulus window exceeded a criterion defined on the interval between the minium and maximium spike count. We then selected the (optimal) criterion that best minimised misses and false alarms. Trials exceeding the optimal criterion were counted as a response. Following Takahashi et al (2016), we then fit logistic functions (equation 14) to the network responses using non-linear least- squares.

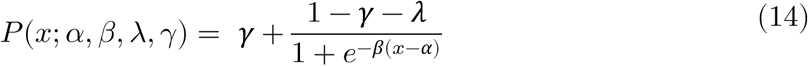

Where *P(x)* is the detection probability (i.e. the probability of the model producing a hit or false alarm), and 𝛼, 𝛽, 𝜆, 𝛾 are free parameters. We used the optimal criterion found in the unperturbed (i.e. control) simulations in the perturbation simulations.

Second, to ensure that our results were not an artefact of the (optimal) criterion we constructed neurometric functions following the procedure described in the supplementary material of Takahashi et al (2016). Specifically, for each stimulus intensity we constructed ROC curves and then computed the AUC (area under the ROC curve) thereby summing over all criterions. To convert the AUC into a quantity comparable to a psychometric function (i.e. so that each neurometric function vairied between 0 and 1), we normalised the AUC values, 𝑃 (𝑟𝑒𝑠𝑝𝑜𝑛𝑠𝑒) = (𝐴𝑈𝐶 − 𝑖𝑛𝑡𝑒𝑟𝑐𝑒𝑝𝑡) ∗ 𝑚𝑎𝑥. The *intercept* was given by the minimium AUC across all conditions, and the 𝑚𝑎𝑥 was given by the maximium AUC across all conditions.

## Supplementary material

### S1. Apical compartment phase plane

**Figure S1.**
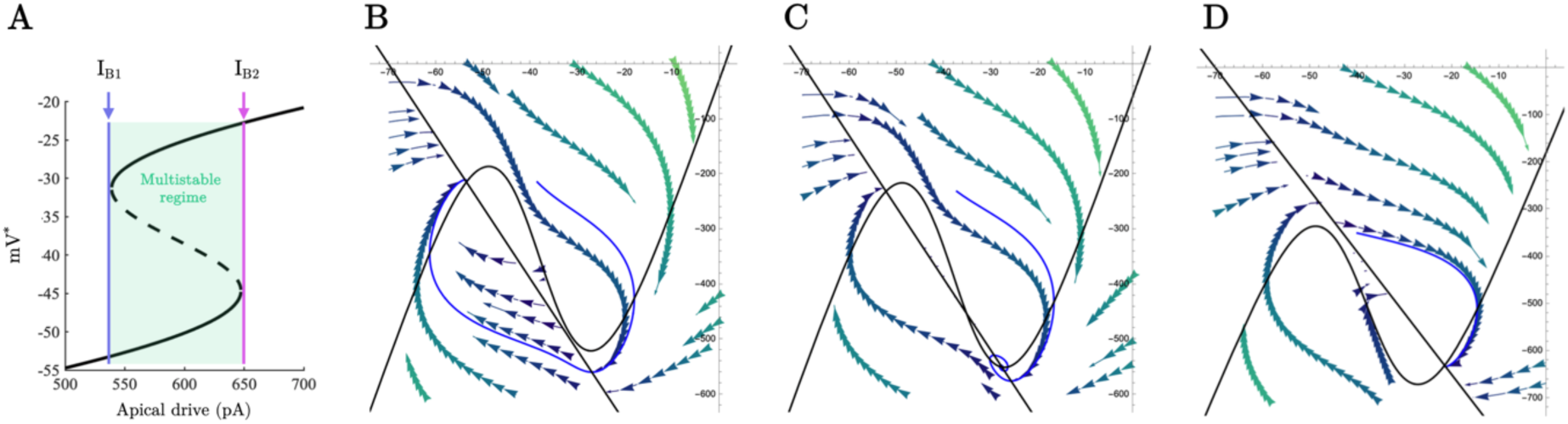
A) Bifurcation diagram of the L5_PT_ apical compartment. The saddle node bifurcation at I_B1_ generates a stable plateau potential which coexists with the resting state of the apical compartment until the model passes through a second saddle node bifurcation at I_B2_ at which point the resting state of the compartment vanishes and the plateau potential becomes globally attracting. B-C) Phase plane representation of the apical compartment showing the nullclines (black) for the following values of the bifurcation parameter; 𝐼_𝑒𝑥𝑡_ < I_B1_, I_B1_ < 𝐼_𝑒𝑥𝑡_< I_B2_, 𝐼_𝑒𝑥𝑡_ >I_B2._

### S2. Sweeping the magnitude of model perturbations

**Figure S2.**
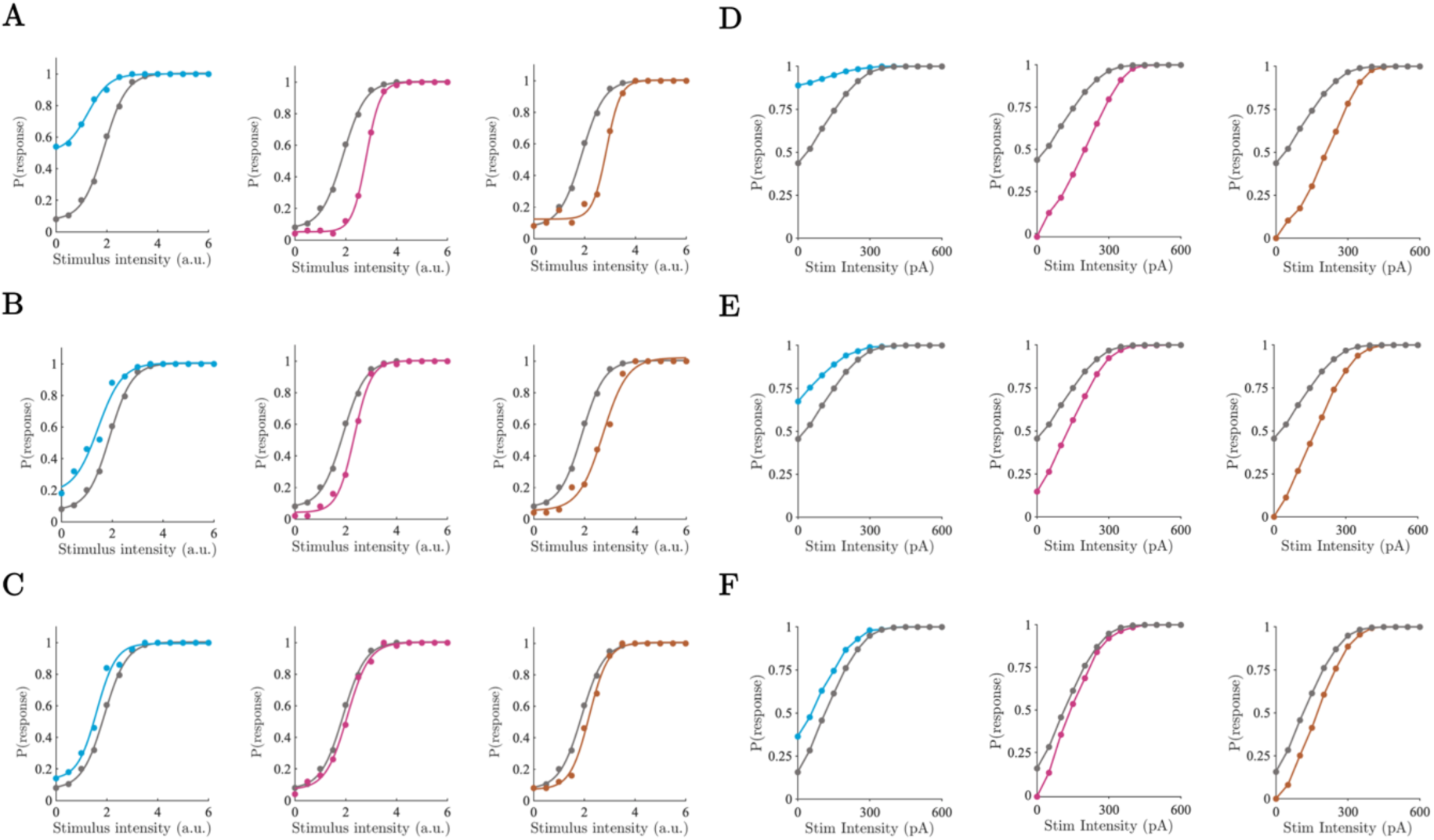
A-C) Psychometric function fit to spiking model output across apical compartment excitation (blue), apical compartment inhibition (pink), and thalamic inhibition (orange), of varying magnitudes; A = 300 pA, B = 200 pA, C = 100 pA. D-F) Same as A-C but for neurometric functions; D = 300 pA, E = 200 pA, F = 100 pA.

### S3. Robustness of rivalry duration across burstiness parameters

**Figure S3.**
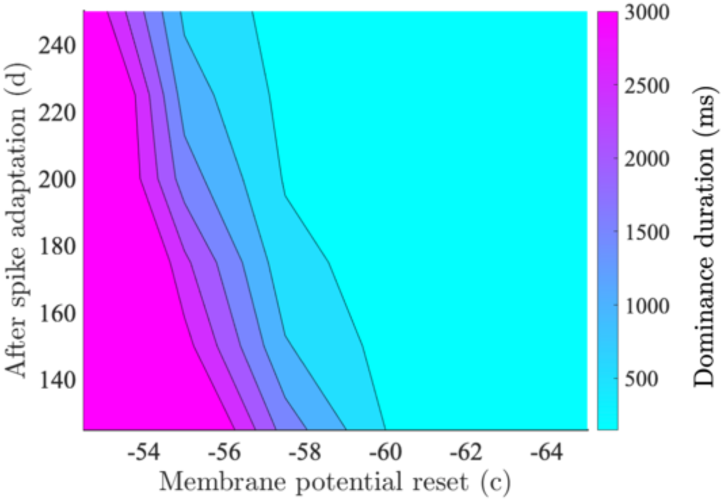
Mean dominance duration as a function of the spike reset parameter values controlling the burstiness of the model L5_PT_ cells.

### S4. Dynamical regime underlying visual rivalry

To interrogate the structure of the dynamical system underlying the stochastic oscillations we made inter-compartment coupling deterministic and drove the model with a constant current and asymmetric initial conditions so that the system converged to a state where one of the populations was dominant whilst the other was suppressed. We reasoned that if the oscillations were driven by stochastic jumps between stable fixed points with basins of attraction modulated by adaptation then in the absence of noise the oscillations should disappear. In contrast, if adaptation exerts a large effect, the oscillations should consist of a stable limit cycle and the model should continue to oscillate in the absence of noise (c.f. Moreno-Bote et al., 2007; Shpiro et al., 2009). In agreement with the stable limit cycle hypothesis in the absence of noise the model continued to oscillate (Fig S4A). To test the stability of the limit cycle we: i) confirmed the existence of an unstable structure inside the limit cycle; and ii) confirmed that perturbations to the limit cycle decayed back to a stable orbit. With symmetric initial conditions the model converged to a state where excitatory activity on each side of the ring was perfectly matched. Perturbations to this state consisting of a 1 ms pulse of constant drive (50 [pA]) to the somatic compartment of a single L5PT neuron caused the orbit to converge to the surrounding limit cycle (Fig S4B). Such perturbations had little effect on already oscillating orbits confirming the stability of the limit cycle (Fig S4C).

**Figure S4.**
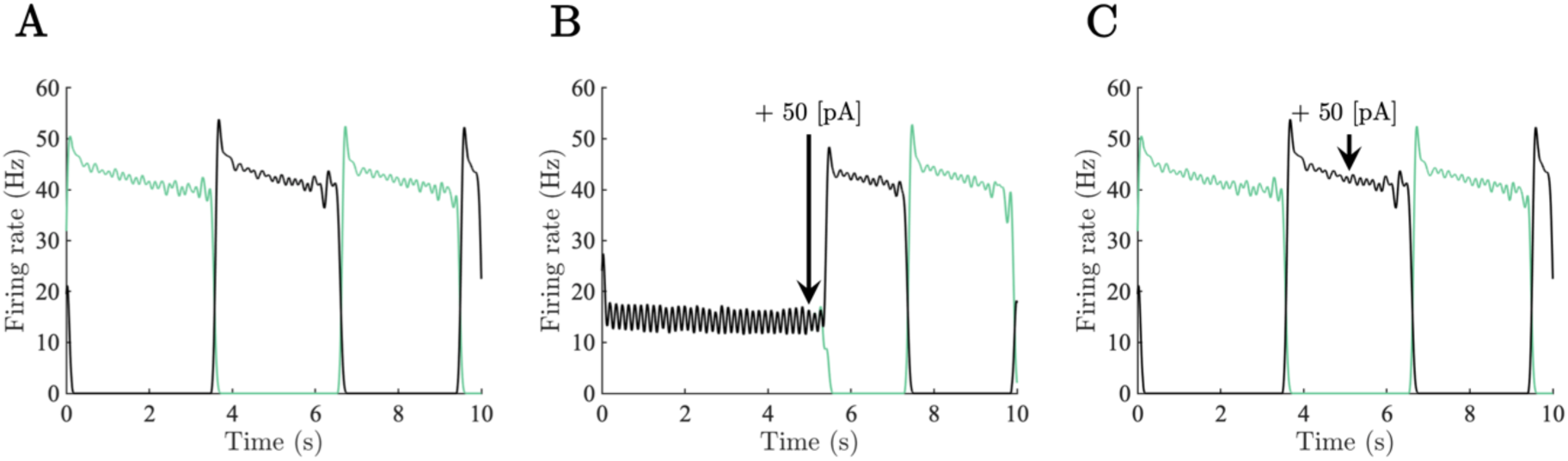
A) Average firing rate of neuronal populations centred on opposite ends of the ring driven by a constant drive with asymmetric initial conditions. B) Average firing rate of neuronal populations simulated with constant drive and symmetric initial conditions. A perturbation was delivered at 𝑡 = 5000 [ms] sending the population orbit to the surrounding stable limit cycle. C) Average firing rate of neuronal populations driven by constant drive with asymmetric initial conditions. Perturbation delivered at 𝑡 = 5000 [ms] had no substantial effect on the already oscillating orbit indicative of a stable limit cycle.

### S5. Key effects of visual rivalry simulations are preserved in scaled-up model

To help guard against possible biases in the results caused by finite size effects or other simplifying assumptions made in the model such as the 50/50 excitatory/inhibitory neuron ratio, or the all-to-all connectivity of the cortical ring we constructed a scaled-up version of the model consisting of 2160 neurons (1600 excitatory, 400 inhibitory, 160 thalamic). The scaled-up model had sparse connectivity (12.5% connection probability), and an 80/20 excitatory/inhibitory neuron ratio (i.e. in line with Dale’s law). Because of the non-linearities in the model, and the reduction in the number of inhibitory neurons, we could not simply rescale the parameters of the original smaller network. Instead, we retuned the connectivity and adaptation parameters using the procedure described in materials and methods. Parameter values of the scaled-up network model are supplied below in table S1.

**Table S1.**
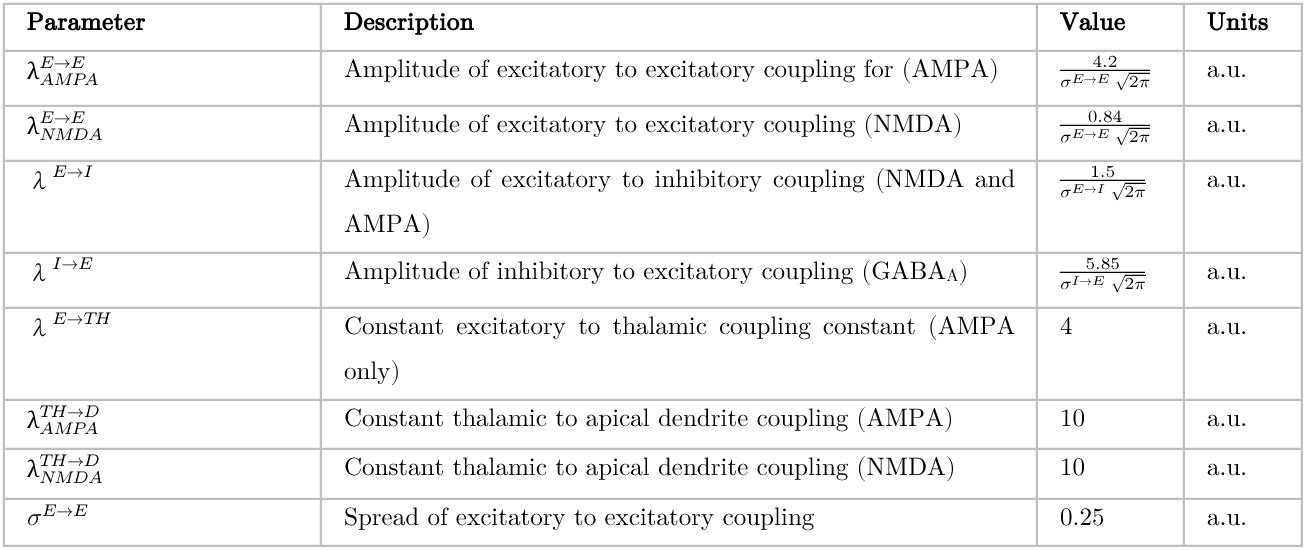
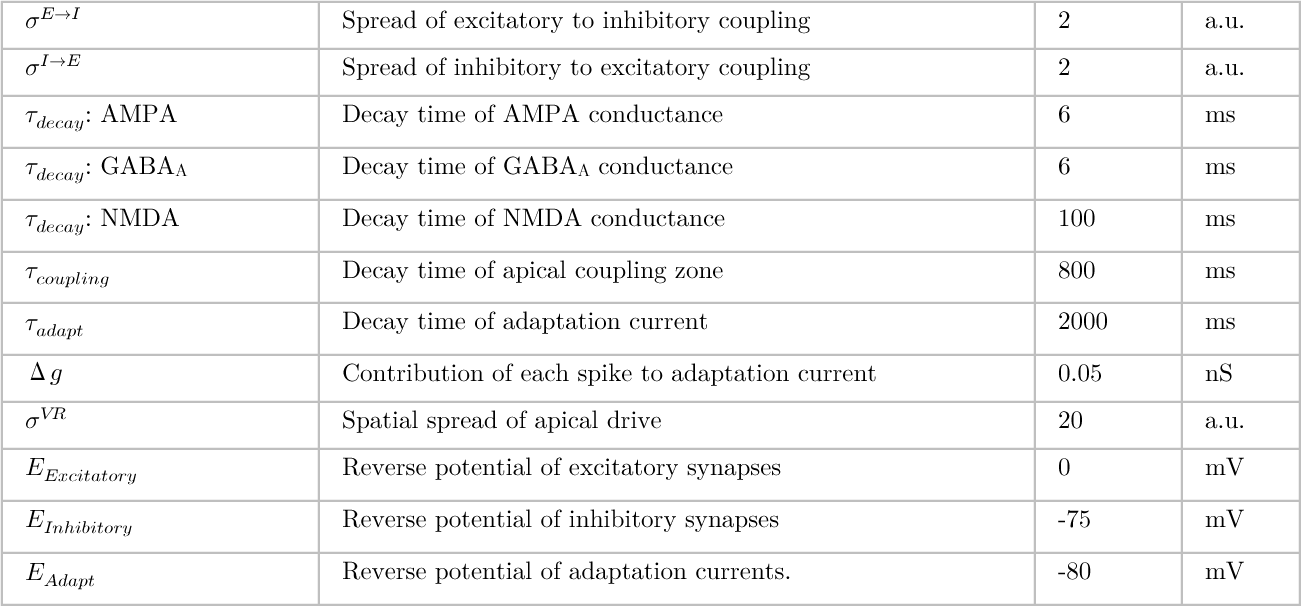
Parameter description, values, and units of the (scaled-up) model components described by equations (5-11).

| Parameter | Description | Value | Units |
| --- | --- | --- | --- |
| $\lambda_{AMPA}^{E \rightarrow E}$ | Amplitude of excitatory to excitatory coupling for (AMPA) | $\frac{4.2}{\sigma^{E \rightarrow E} \sqrt{2\pi}}$ | a.u. |
| $\lambda_{NMDA}^{E \rightarrow E}$ | Amplitude of excitatory to excitatory coupling (NMDA) | $\frac{0.84}{\sigma^{E \rightarrow E} \sqrt{2\pi}}$ | a.u. |
| $\lambda^{E \rightarrow I}$ | Amplitude of excitatory to inhibitory coupling (NMDA and AMPA) | $\frac{1.5}{\sigma^{E \rightarrow I} \sqrt{2\pi}}$ | a.u. |
| $\lambda^{I \rightarrow E}$ | Amplitude of inhibitory to excitatory coupling (GABA<sub>A</sub>) | $\frac{5.85}{\sigma^{I \rightarrow E} \sqrt{2\pi}}$ | a.u. |
| $\lambda^{E \rightarrow TH}$ | Constant excitatory to thalamic coupling constant (AMPA only) | 4 | a.u. |
| $\lambda_{AMPA}^{TH \rightarrow D}$ | Constant thalamic to apical dendrite coupling (AMPA) | 10 | a.u. |
| $\lambda_{NMDA}^{TH \rightarrow D}$ | Constant thalamic to apical dendrite coupling (NMDA) | 10 | a.u. |
| $\sigma^{E \rightarrow E}$ | Spread of excitatory to excitatory coupling | 0.25 | a.u. |
| $\sigma^{E \rightarrow I}$ | Spread of excitatory to inhibitory coupling | 2 | a.u. |
| $\sigma^{I \rightarrow E}$ | Spread of inhibitory to excitatory coupling | 2 | a.u. |
| $\tau_{decay}$: AMPA | Decay time of AMPA conductance | 6 | ms |
| $\tau_{decay}$: GABA<sub>A</sub> | Decay time of GABA<sub>A</sub> conductance | 6 | ms |
| $\tau_{decay}$: NMDA | Decay time of NMDA conductance | 100 | ms |
| $\tau_{coupling}$ | Decay time of apical coupling zone | 800 | ms |
| $\tau_{adapt}$ | Decay time of adaptation current | 2000 | ms |
| $\Delta g$ | Contribution of each spike to adaptation current | 0.05 | nS |
| $\sigma^{VR}$ | Spatial spread of apical drive | 20 | a.u. |
| $E_{Excitatory}$ | Reverse potential of excitatory synapses | 0 | mV |
| $E_{Inhibitory}$ | Reverse potential of inhibitory synapses | -75 | mV |
| $E_{Adapt}$ | Reverse potential of adaptation currents. | -80 | mV |

In line with the behaviour of the small network model reported in the main text, the large network model (Fig S5A) generated a Gamma distribution of dominance durations (Fig S5B), and was consistent with Levelt’s second (Fig S5C) and fourth (Fig S5D) propositions supporting the robustness of the burst-dependent mechanism of perceptual dominance put forward in the paper. All simulations run in the scaled up network lasted for 20 [s] and results were averaged over 20 random seeds.

**Figure S5.**
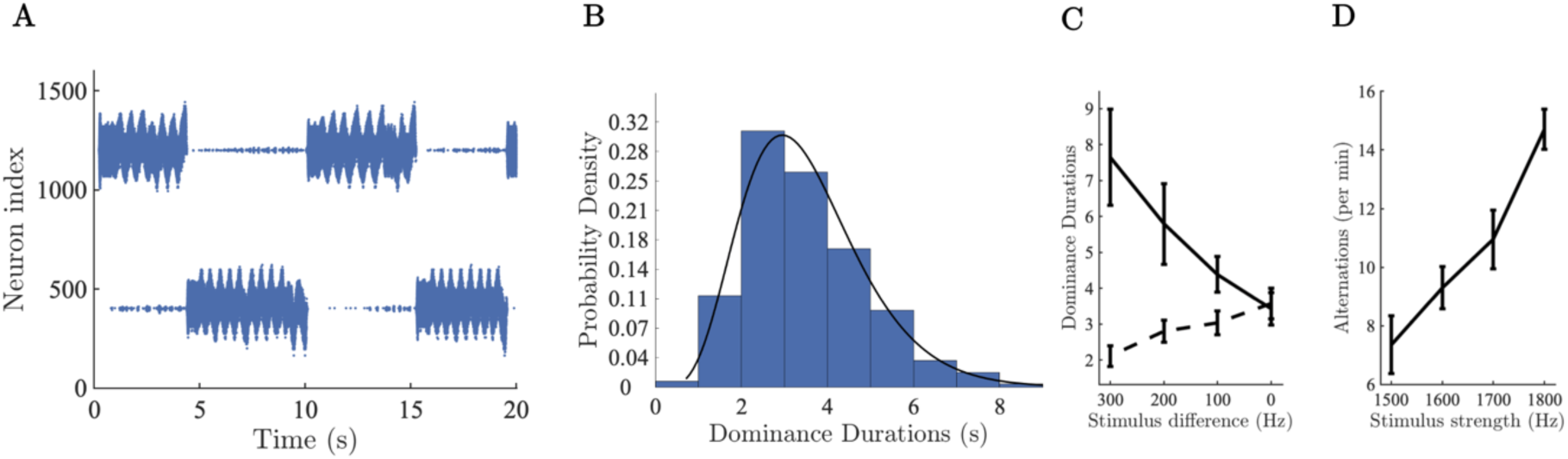
A) Raster plots of somatic spikes from the scaled-up population of L5_PT_ cells B) Histogram of dominance durations, black line shows the fit of a Gamma distribution with parameters estimated via MLE (𝛼 = 6.2, 𝜃 = 0.56). C) Simulation confirming Levelt’s second proposition in scaled-up model. Dashed line shows the dominance duration of the population receiving the decreasing external drive, solid line shows dominance duration of population receiving a fixed drive. D) Simulation of Levelt’s fourth proposition in scaled-up model.

### S6. In silico electrophysiology supplemental figures

**Figure S6.**
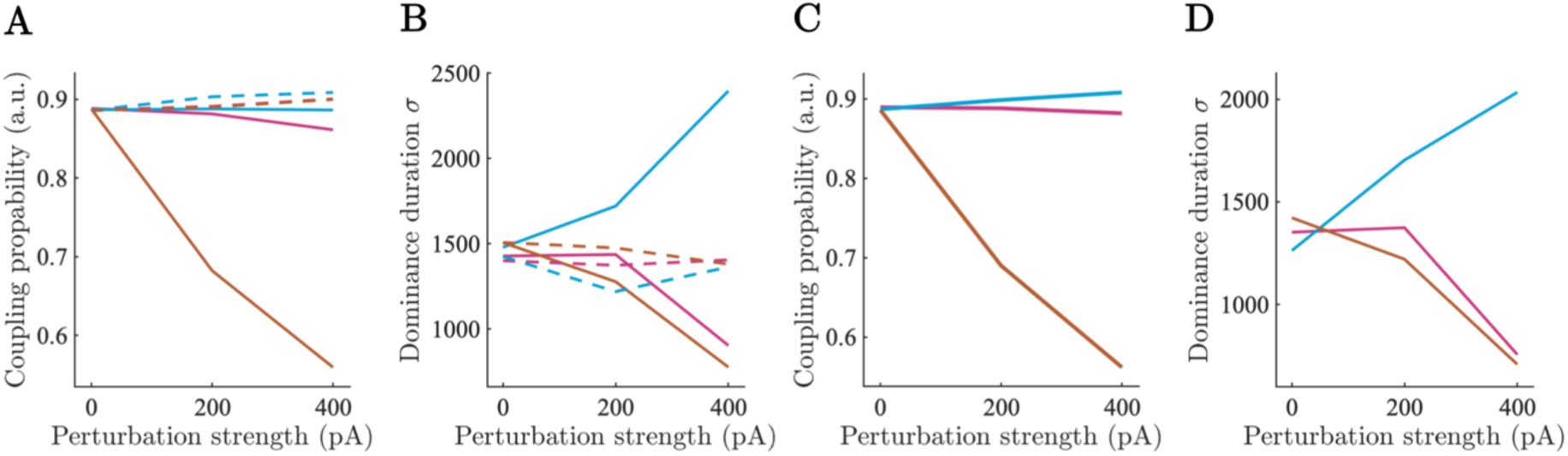
A) Inter-compartment coupling probability under asymmetric perturbation for perturbed (solid) and unperturbed (dashed) populations as a function of perturbation strength for each perturbation type (colours same as main text). B) Dominance duration standard deviation under asymmetric perturbation as a function of the strength of each perturbation type. C) Inter-compartment coupling probability under symmetric perturbations. D) Dominance duration standard deviation under symmetric perturbation as a function of perturbation strength for each perturbation type.

## Code and Data Availability

Complete code necessary to reproduce the simulations reported in the paper can be found at https://github.com/cjwhyte/LVPA.

## Acknowledgements

We are grateful to Michael Breakspear and Bryan Paton for insightful discussion on the initial development of the model. The Sydney Systems Neuroscience and Complexity group, David Alais, Matt Davidson, Jacob Coorey, and Hugh Wilson provided discerning comments and advice that helped to shape the presentation and structure of the manuscript. Finally, we thank Christopher Conn for high-performance computing assistance.

## References

1. Alais, D., & Blake, R. (2005). Binocular Rivalry. MIT Press.

2. Albantakis, L., Barbosa, L., Findlay, G., Grasso, M., Haun, A. M., Marshall, W., Mayner, W. G. P., Zaeemzadeh, A., Boly, M., Juel, B. E., Sasai, S., Fujii, K., David, I., Hendren, J., Lang, J. P., & Tononi, G. (2023). Integrated information theory (IIT) 4.0: Formulating the properties of phenomenal existence in physical terms. PLOS Computational Biology, 19(10), e1011465. 10.1371/journal.pcbi.1011465

3. Aru, J., Suzuki, M., & Larkum, M. E. (2020). Cellular Mechanisms of Conscious Processing. Trends in Cognitive Sciences, 24(10), 814–825. 10.1016/j.tics.2020.07.006

4. Aru, J., Suzuki, M., Rutiku, R., Larkum, M. E., & Bachmann, T. (2019). Coupling the State and Contents of Consciousness. Frontiers in Systems Neuroscience, 13, 43. 10.3389/fnsys.2019.00043

5. Bachmann, T., Suzuki, M., & Aru, J. (2020). Dendritic integration theory: A thalamo-cortical theory of state and content of consciousness. Philosophy and the Mind Sciences, 1(II), Article II. 10.33735/phimisci.2020.II.52

6. Benitez, F., Pennartz, C., & Senn, W. (2023). The conductor model of consciousness, our neuromorphic twins, and the human-AI deal. 10.31234/osf.io/gbzd6

7. Blake, R., & Tong, F. (2008). Binocular rivalry. Scholarpedia, 3(12), 1578. 10.4249/scholarpedia.1578

8. Bogatova, D., Smirnakis, S. M., & Palagina, G. (2024). Tug-of-Peace: Visual Rivalry and Atypical Visual Motion Processing in MECP2 Duplication Syndrome of Autism. Eneuro, 11(1), ENEURO.0102-23.2023. 10.1523/ENEURO.0102-23.2023

9. Bonneh, Y. S., Donner, T. H., Cooperman, A., Heeger, D. J., & Sagi, D. (2014). Motion-Induced Blindness and Troxler Fading: Common and Different Mechanisms. PLoS ONE, 9(3), e92894. 10.1371/journal.pone.0092894

10. Brascamp, J. W., Klink, P. C., & Levelt, W. J. M. (2015). The ‘laws’ of binocular rivalry: 50 years of Levelt’s propositions. Vision Research, 109, 20–37. 10.1016/j.visres.2015.02.019

11. Brascamp, J. W., van Ee, R., Noest, A. J., Jacobs, R. H. A. H., & van den Berg, A. V. (2006). The time course of binocular rivalry reveals a fundamental role of noise. Journal of Vision, 6(11), 8–8. 10.1167/6.11.8

12. Britten, K., Shadlen, M., Newsome, W., & Movshon, J. (1992). The analysis of visual motion: A comparison of neuronal and psychophysical performance. The Journal of Neuroscience, 12(12), 4745–4765. 10.1523/JNEUROSCI.12-12-04745.1992

13. Buckthought, A., Kim, J., & Wilson, H. R. (2008). Hysteresis effects in stereopsis and binocular rivalry. Vision Research, 48(6), 819–830. 10.1016/j.visres.2007.12.013

14. Carmel, D., Arcaro, M., Kastner, S., & Hasson, U. (2010). How to Create and Use Binocular Rivalry. Journal of Visualized Experiments, 45, 2030. 10.3791/2030

15. Ciceri, S., Lohuis, M. N., Rottschäfer, V., Pennartz, C. M., Avitabile, D., van Gaal, S., & Olcese, U. (2024). The neural and computational architecture of feedback dynamics in mouse cortex during stimulus report (p. 2023.07.19.549692). bioRxiv. 10.1101/2023.07.19.549692

16. Cortes, N., Ladret, H. J., Abbas-Farishta, R., & Casanova, C. (2023). The pulvinar as a hub of visual processing and cortical integration. *Trends in Neurosciences*, S0166223623002709. 10.1016/j.tins.2023.11.008

17. Dayan, P., & Abbott, L. F. (2005). Theoretical Neuroscience: Computational and Mathematical Modeling of Neural Systems. MIT Press.

18. Destexhe, A. (2009). Self-sustained asynchronous irregular states and Up–Down states in thalamic, cortical and thalamocortical networks of nonlinear integrate-and-fire neurons. Journal of Computational Neuroscience, 27(3), 493–506. 10.1007/s10827-009-0164-4

19. Douglas, R. J., & Martin, K. A. C. (2004). Neuronal Circuits of the Neocortex. Annual Review of Neuroscience, 27(1), 419–451. 10.1146/annurev.neuro.27.070203.144152

20. Dwarakanath, A., Kapoor, V., Werner, J., Safavi, S., Fedorov, L. A., Logothetis, N. K., & Panagiotaropoulos, T. I. (2020). *Prefrontal state fluctuations control access to consciousness* [Preprint]. Neuroscience. 10.1101/2020.01.29.924928

21. Gale, S. D., Strawder, C., Bennett, C., Mihalas, S., Koch, C., & Olsen, S. R. (2024). Backward masking in mice requires visual cortex. Nature Neuroscience, 27(1), 129–136. 10.1038/s41593-023-01488-0

22. Gerstner, W., Kistler, W. M., Naud, R., & Paninski, L. (2014). Neuronal Dynamics: From Single Neurons to Networks and Models of Cognition (1st ed.). Cambridge University Press. 10.1017/CBO9781107447615

23. Greget, R., Pernot, F., Bouteiller, J.-M. C., Ghaderi, V., Allam, S., Keller, A. F., Ambert, N., Legendre, A., Sarmis, M., Haeberle, O., Faupel, M., Bischoff, S., Berger, T. W., & Baudry, M. (2011). Simulation of Postsynaptic Glutamate Receptors Reveals Critical Features of Glutamatergic Transmission. PLOS ONE, 6(12), e28380. 10.1371/journal.pone.0028380

24. Grossberg, S., Yazdanbakhsh, A., Cao, Y., & Swaminathan, G. (2008). How does binocular rivalry emerge from cortical mechanisms of 3-D vision? Vision Research, 48(21), 2232–2250. 10.1016/j.visres.2008.06.024

25. Harris, K. D., & Shepherd, G. M. G. (2015). The neocortical circuit: Themes and variations. Nature Neuroscience, 18(2), 170–181. 10.1038/nn.3917

26. He, B. J. (2023). Next frontiers in consciousness research. Neuron, 111(20), 3150– 3153. 10.1016/j.neuron.2023.09.042

27. Hupé, J.-M., Signorelli, C. M., & Alais, D. (2019). Two paradigms of bistable plaid motion reveal independent mutual inhibition processes. Journal of Vision, 19(4), 5. 10.1167/19.4.5

28. Izhikevich, E. M. (2003). Simple model of spiking neurons. IEEE Transactions on Neural Networks, 14(6), 1569–1572. IEEE Transactions on Neural Networks. 10.1109/TNN.2003.820440

29. Izhikevich, E. M. (2004). Which model to use for cortical spiking neurons? IEEE Transactions on Neural Networks, 15(5), 1063–1070. IEEE Transactions on Neural Networks. 10.1109/TNN.2004.832719

30. Izhikevich, E. M. (2006). Dynamical Systems in Neuroscience: The Geometry of Excitability and Bursting. The MIT Press. 10.7551/mitpress/2526.001.0001

31. Izhikevich, E. M. (2010). Hybrid spiking models. *Philosophical Transactions of the Royal Society A: Mathematical*, Physical and Engineering Sciences, 368(1930), 5061–5070. 10.1098/rsta.2010.0130

32. Izhikevich, E. M., & Edelman, G. M. (2008). Large-scale model of mammalian thalamocortical systems. Proceedings of the National Academy of Sciences, 105(9), 3593–3598. 10.1073/pnas.0712231105

33. Jaramillo, J., Mejias, J. F., & Wang, X.-J. (2019). Engagement of Pulvino-cortical Feedforward and Feedback Pathways in Cognitive Computations. Neuron, 101(2), 321–336.e9. 10.1016/j.neuron.2018.11.023

34. Kalmbach, B. E., Hodge, R. D., Jorstad, N. L., Owen, S., De Frates, R., Yanny, A. M., Dalley, R., Mallory, M., Graybuck, L. T., Radaelli, C., Keene, C. D., Gwinn, R. P., Silbergeld, D. L., Cobbs, C., Ojemann, J. G., Ko, A. L., Patel, A. P., Ellenbogen, R. G., Bakken, T. E., … Ting, J. T. (2021). Signature morpho-electric, transcriptomic, and dendritic properties of human layer 5 neocortical pyramidal neurons. Neuron, 109(18), 2914–2927.e5. 10.1016/j.neuron.2021.08.030

35. Kapoor, V., Dwarakanath, A., Safavi, S., Werner, J., Besserve, M., Panagiotaropoulos, T. I., & Logothetis, N. K. (2020). *Decoding the contents of consciousness from prefrontal ensembles* [Preprint]. Neuroscience. 10.1101/2020.01.28.921841

36. Klatzmann, U., Froudist-Walsh, S., Bliss, D. P., Theodoni, P., Mejias, J. F., Niu, M., Rapan, L., Palomero-Gallagher, N., Sergent, C., Dehaene, S., & Wang, X.-J. (2022). *A connectome-based model of conscious access in monkey cortex* [Preprint]. Neuroscience. 10.1101/2022.02.20.481230

37. Laing, C. R., & Chow, C. C. (2002). A Spiking Neuron Model for Binocular Rivalry. Journal of Computational Neuroscience, 12(1), 39–53. 10.1023/A:1014942129705

38. Laing, C. R., Frewen, T., & Kevrekidis, I. G. (2010). Reduced models for binocular rivalry. Journal of Computational Neuroscience, 28(3), 459–476. 10.1007/s10827-010-0227-6

39. Lamme, V. A. F. (2006). Towards a true neural stance on consciousness. Trends in Cognitive Sciences, 10(11), 494–501. 10.1016/j.tics.2006.09.001

40. Larkum, M. E. (2004). Top-down Dendritic Input Increases the Gain of Layer 5 Pyramidal Neurons. Cerebral Cortex, 14(10), 1059–1070. 10.1093/cercor/bhh065

41. Larkum, M. E. (2022). Are Dendrites Conceptually Useful? Neuroscience, 489, 4–14. 10.1016/j.neuroscience.2022.03.008

42. Larkum, M. E., Nevian, T., Sandler, M., Polsky, A., & Schiller, J. (2009). Synaptic Integration in Tuft Dendrites of Layer 5 Pyramidal Neurons: A New Unifying Principle. Science, 325(5941), 756–760. 10.1126/science.1171958

43. Lau, H. C. (2007). A higher order Bayesian decision theory of consciousness. In Progress in Brain Research (Vol. 168, pp. 35–48). Elsevier. 10.1016/S0079-6123(07)68004-2

44. Levelt, W. J. M. (1965). On Binocular Rivalry [Institute for Perception]. https://www.mpi.nl/world/materials/publications/levelt/Levelt_Binocular_Rivalry_1965.pdf

45. Levelt, W. J. M. (1967). Note on the Distribution of Dominance Times in Binocular Rivalry. British Journal of Psychology, 58(1–2), 143–145. 10.1111/j.2044-8295.1967.tb01068.x

46. Li, H.-H., Rankin, J., Rinzel, J., Carrasco, M., & Heeger, D. J. (2017). Attention model of binocular rivalry. Proceedings of the National Academy of Sciences, 114(30). 10.1073/pnas.1620475114

47. Logothetis, N. K., & Leopold, D. A. (1996). *What is rivalling during binocular rivalry? 380*.

48. Marvan, T., Polák, M., Bachmann, T., & Phillips, W. A. (2021). Apical amplification—A cellular mechanism of conscious perception? Neuroscience of Consciousness, 2021(2), niab036. 10.1093/nc/niab036

49. Mashour, G. A., Roelfsema, P., Changeux, J.-P., & Dehaene, S. (2020). Conscious Processing and the Global Neuronal Workspace Hypothesis. Neuron, 105(5), 776–798. 10.1016/j.neuron.2020.01.026

50. McCormick, D. A., & Williamson, A. (1989). Convergence and divergence of neurotransmitter action in human cerebral cortex. Proceedings of the National Academy of Sciences, 86(20), 8098–8102. 10.1073/pnas.86.20.8098

51. Mease, R. A., & Gonzalez, A. J. (2021). Corticothalamic Pathways From Layer 5: Emerging Roles in Computation and Pathology. Frontiers in Neural Circuits, 15, 730211. 10.3389/fncir.2021.730211

52. Meng, M., & Tong, F. (2004). Can attention selectively bias bistable perception? Differences between binocular rivalry and ambiguous figures. Journal of Vision, 4(7), 2. 10.1167/4.7.2

53. Moreno-Bote, R., Rinzel, J., & Rubin, N. (2007). Noise-Induced Alternations in an Attractor Network Model of Perceptual Bistability. Journal of Neurophysiology, 98(3), 1125–1139. 10.1152/jn.00116.2007

54. Müller, E. J., Munn, B. R., Redinbaugh, M. J., Lizier, J., Breakspear, M., Saalmann, Y. B., & Shine, J. M. (2023). The non-specific matrix thalamus facilitates the cortical information processing modes relevant for conscious awareness. Cell Reports, 42(8), 112844. 10.1016/j.celrep.2023.112844

55. Müller, E. J., Munn, B. R., & Shine, J. M. (2020). Diffuse neural coupling mediates complex network dynamics through the formation of quasi-critical brain states. Nature Communications, 11(1), 6337. 10.1038/s41467-020-19716-7

56. Munn, B. R., Müller, E. J., Aru, J., Whyte, C. J., Gidon, A., Larkum, M. E., & Shine, J. M. (2023). A thalamocortical substrate for integrated information via critical synchronous bursting. Proceedings of the National Academy of Sciences, 120(46), e2308670120. 10.1073/pnas.2308670120

57. Munn, B. R., Müller, E. J., Medel, V., Naismith, S. L., Lizier, J. T., Sanders, R. D., & Shine, J. M. (2023). Neuronal connected burst cascades bridge macroscale adaptive signatures across arousal states. Nature Communications, 14(1), Article 1. 10.1038/s41467-023-42465-2

58. Naka, A., & Adesnik, H. (2016). Inhibitory Circuits in Cortical Layer 5. Frontiers in Neural Circuits, 10. 10.3389/fncir.2016.00035

59. Naud, R., Bathellier, B., & Gerstner, W. (2014). Spike-timing prediction in cortical neurons with active dendrites. Frontiers in Computational Neuroscience, 8. 10.3389/fncom.2014.00090

60. Naud, R., & Sprekeler, H. (2018). Sparse bursts optimize information transmission in a multiplexed neural code. Proceedings of the National Academy of Sciences, 115(27). 10.1073/pnas.1720995115

61. Oude Lohuis, M. N., Pie, J. L., Marchesi, P., Montijn, J. S., de Kock, C. P. J., Pennartz, C. M. A., & Olcese, U. (2022). Multisensory task demands temporally extend the causal requirement for visual cortex in perception. Nature Communications, 13(1), 2864. 10.1038/s41467-022-30600-4

62. Overgaard, M. (2015). Behavioural Methods in Consciousness Research. Oxford University Press.

63. Palagina, G., Meyer, J. F., & Smirnakis, S. M. (2017). Complex Visual Motion Representation in Mouse Area V1. The Journal of Neuroscience, 37(1), 164–183. 10.1523/JNEUROSCI.0997-16.2017

64. Phillips, W. A., Larkum, M. E., Harley, C. W., & Silverstein, S. M. (2016). The effects of arousal on apical amplification and conscious state. Neuroscience of Consciousness, 2016(1), niw015. 10.1093/nc/niw015

65. Pitts, M. A., Metzler, S., & Hillyard, S. A. (2014). Isolating neural correlates of conscious perception from neural correlates of reporting one’s perception. Frontiers in Psychology, 5. 10.3389/fpsyg.2014.01078

66. Poort, J., & Meyer, A. F. (2021). Vision: Depth perception in climbing mice. Current Biology, 31(10), R486–R488. 10.1016/j.cub.2021.03.066

67. Pujol, C. F., Blundon, E. G., & Dykstra, A. R. (2023). Laminar specificity of the auditory perceptual awareness negativity: A biophysical modeling study. PLOS Computational Biology, 19(6), e1011003. 10.1371/journal.pcbi.1011003

68. Qian, C., Chen, Z., de Hollander, G., Knapen, T., Zhang, Z., He, S., & Zhang, P. (2023). Hierarchical and fine-scale mechanisms of binocular rivalry for conscious perception (p. 2023.02.11.528110). bioRxiv. 10.1101/2023.02.11.528110

69. Safavi, S., & Dayan, P. (2022). Multistability, perceptual value, and internal foraging. Neuron, S0896627322006699. 10.1016/j.neuron.2022.07.024

70. Seo, J., Kim, D.-J., Choi, S.-H., Kim, H., & Min, B.-K. (2022). The thalamocortical inhibitory network controls human conscious perception. NeuroImage, 264, 119748. 10.1016/j.neuroimage.2022.119748

71. Sergent, C., Baillet, S., & Dehaene, S. (2005). Timing of the brain events underlying access to consciousness during the attentional blink. Nature Neuroscience, 8(10), 1391–1400. 10.1038/nn1549

72. Shai, A. S., Anastassiou, C. A., Larkum, M. E., & Koch, C. (2015). Physiology of Layer 5 Pyramidal Neurons in Mouse Primary Visual Cortex: Coincidence Detection through Bursting. PLOS Computational Biology, 11(3), e1004090. 10.1371/journal.pcbi.1004090

73. Shepherd, G. M. G., & Yamawaki, N. (2021). Untangling the cortico-thalamo- cortical loop: Cellular pieces of a knotty circuit puzzle. Nature Reviews Neuroscience, 22(7), 389–406. 10.1038/s41583-021-00459-3

74. Shpiro, A., Curtu, R., Rinzel, J., & Rubin, N. (2007). Dynamical Characteristics Common to Neuronal Competition Models. Journal of Neurophysiology, 97(1), 462–473. 10.1152/jn.00604.2006

75. Shpiro, A., Moreno-Bote, R., Rubin, N., & Rinzel, J. (2009). Balance between noise and adaptation in competition models of perceptual bistability. Journal of Computational Neuroscience, 27(1), 37–54. 10.1007/s10827-008-0125-3

76. Storm, J. F., Klink, P. C., Aru, J., Senn, W., Goebel, R., Pigorini, A., Avanzini, P., Vanduffel, W., Roelfsema, P. R., Massimini, M., Larkum, M. E., & Pennartz, C. M. A. (2024). An integrative, multiscale view on neural theories of consciousness. Neuron, 112(10), 1531–1552. 10.1016/j.neuron.2024.02.004

77. Strogatz, S. (2018). Nonlinear dynamics and chaos: With applications to physics, biology, chemistry, and engineering (Second edition). CRC Press Taylor & Francis Group.

78. Suzuki, M., & Larkum, M. E. (2020). General Anesthesia Decouples Cortical Pyramidal Neurons. Cell, 180(4), 666–676.e13. 10.1016/j.cell.2020.01.024

79. Tahvili, F., & Destexhe, A. (2023). *A mean-field model of gamma-frequency oscillations in networks of excitatory and inhibitory neurons* [Preprint]. Neuroscience. 10.1101/2023.11.20.567709

80. Takahashi, N., Ebner, C., Sigl-Glöckner, J., Moberg, S., Nierwetberg, S., & Larkum, M. E. (2020). Active dendritic currents gate descending cortical outputs in perception. Nature Neuroscience, 23(10), 1277–1285. 10.1038/s41593-020-0677-8

81. Takahashi, N., Oertner, T. G., Hegemann, P., & Larkum, M. E. (2016c). Active cortical dendrites modulate perception. Science, 354(6319), 1587–1590. 10.1126/science.aah6066

82. Tong, F., Meng, M., & Blake, R. (2006). Neural bases of binocular rivalry. Trends in Cognitive Sciences, 10(11), 502–511. 10.1016/j.tics.2006.09.003

83. Trautmann, E. M., Hesse, J. K., Stine, G. M., Xia, R., Zhu, S., O’Shea, D. J., Karsh, B., Colonell, J., Lanfranchi, F. F., Vyas, S., Zimnik, A., Steinmann, N. A., Wagenaar, D. A., Andrei, A., Lopez, C. M., O’Callaghan, J., Putzeys, J., Raducanu, B. C., Welkenhuysen, M., … Harris, T. (2023). *Large-scale high-density brain-wide neural recording in nonhuman primates* [Preprint]. Neuroscience. 10.1101/2023.02.01.526664

84. Wang, Z., Dai, W., & McLaughlin, D. W. (2020). Ring models of binocular rivalry and fusion. Journal of Computational Neuroscience, 48(2), 193–211. 10.1007/s10827-020-00744-7

85. Whyte, C. J., Redinbaugh, M. J., Shine, J. M., & Saalmann, Y. B. (2024). Thalamic contributions to the state and contents of consciousness. Neuron, 112(10), 1611–1625. 10.1016/j.neuron.2024.04.019

86. Wilke, M., Mueller, K.-M., & Leopold, D. A. (2009). Neural activity in the visual thalamus reflects perceptual suppression. Proceedings of the National Academy of Sciences, 106(23), 9465–9470. 10.1073/pnas.0900714106

87. Wilson, H. R. (2003). Computational evidence for a rivalry hierarchy in vision. Proceedings of the National Academy of Sciences, 100(24), 14499–14503. 10.1073/pnas.2333622100

88. Wilson, H. R. (2007). Minimal physiological conditions for binocular rivalry and rivalry memory. Vision Research, 47(21), 2741–2750. 10.1016/j.visres.2007.07.007

89. Wilson, H. R. (2017). Binocular contrast, stereopsis, and rivalry: Toward a dynamical synthesis. Vision Research, 140, 89–95. 10.1016/j.visres.2017.07.016

90. Wilson, H. R., & Cowan, J. D. (2021). Evolution of the Wilson–Cowan equations. Biological Cybernetics, 115(6), 643–653. 10.1007/s00422-021-00912-7

91. Zerlaut, Y., Chemla, S., Chavane, F., & Destexhe, A. (2018). Modeling mesoscopic cortical dynamics using a mean-field model of conductance-based networks of adaptive exponential integrate-and-fire neurons. Journal of Computational Neuroscience, 44(1), 45–61. 10.1007/s10827-017-0668-2

